# Initiation of B-type starch granules in wheat endosperm requires the plastidial α-glucan phosphorylase PHS1

**DOI:** 10.1101/2023.06.01.543270

**Authors:** Nitin Uttam Kamble, Farrukh Makhamadnojov, Brendan Fahy, Carlo Martins, Gerhard Saalbach, David Seung

## Abstract

PHS1 is a plastidial α-glucan phosphorylase that can elongate and degrade maltooligosaccharides (MOS), but its exact physiological role in plants is poorly understood. Here, we discover a specialised role of PHS1 in establishing the unique bimodal characteristic of starch granules in the wheat endosperm. Wheat endosperm contains large A-type granules that initiate at early grain development, and small B-type granules that initiate in later grain development. We demonstrate that PHS1 interacts with BGC1 – a carbohydrate-binding protein essential for normal B-type granule initiation. Mutants of tetraploid durum wheat deficient in all homeologs of PHS1 had normal A-type granules, but fewer and larger B-type granules. Grain size and starch content were not affected by the mutations. Further, by assessing granule numbers during grain development in the *phs1* mutant, and using a double mutant defective in both PHS1 and BGC1, we demonstrate that PHS1 is exclusively involved in B-type granule initiation. The total starch content and number of starch granules per chloroplast in leaves were not affected by loss of PHS1, suggesting that its role in granule initiation in wheat is limited to the endosperm. We therefore propose that the initiation of A- and B-type granules occur via distinct biochemical mechanisms, where PHS1 plays an exclusive role in B-type granule initiation.

## Introduction

Starch, the major storage carbohydrate in plants, is synthesised in plastids as semi-crystalline granules composed of two glucose polymers (glucans) - amylopectin and amylose. Recent work has revealed new insights into molecular mechanisms that initiate (or prime) the formation of granules, and control the number, size and shape of granules in plastids (Seung and Smith, 2019; Mérida and Fettke, 2021). However, there is huge variation in the spatiotemporal patterns of granule initiation between different species and different organs (Seung and Smith, 2019; Chen et al., 2021). The endosperm of wheat (and other Triticeae) has two distinct types of granules: large discoid A-type granules and small spherical B-type granules (Langeveld et al., 2000; Chia et al., 2020). A single A-type granule is initiated in each amyloplast during early grain development, and numerous B-type granules are initiated 10-15 days after the B-type granules, at least partially in amyloplast stromules (Parker, 1985; Langeveld et al., 2000; Esch et al., 2023). The mechanisms underpinning this unique spatiotemporal pattern of A- and B-type granule formation are poorly understood. Understanding such mechanisms is also of industrial significance, since granule size distributions affect grain quality for bread and pasta making (Soh et al., 2006; Park et al., 2009), and brewing (Bathgate and Palmer, 1973).

It is not known whether A- and B-type granule initiations occur through similar or distinct biochemical mechanisms. Both STARCH SYNTHASE 4 (SS4) and B-GRANULE CONTENT1 (BGC1) are required for proper A-type granule initiation in wheat. Loss of either protein results in supernumerary granule initiations in most amyloplasts during early grain development, which later fuse to form ‘compound’ starch granules (Chia et al., 2020; Hawkins et al., 2021). However, BGC1 also plays an important role during B-type granule initiation. While SS4 expression levels peak during early grain development, BGC1 expression is highest during later grain development, when B-type granules are initiating (Chen et al., 2022). B-type granule formation can be almost completely eliminated by reducing BGC1 gene dosage: by introducing loss-of-function mutations in two of three homoeologs of BGC1 in hexaploid wheat, or by combining a loss-of-function mutation in one homeolog with a hypomorphic missense mutation in the other homeolog in tetraploid durum wheat (Chia et al., 2020).

The mechanism by which BGC1 acts in B-type granule initiation is not known. BGC1, along with FLO6 in barley and rice, are all orthologs of Arabidopsis PTST2 and thus belong to the PROTEIN TARGETING TO STARCH (PTST) family. These proteins have a Carbohydrate Binding Module (CBM48) that can bind soluble maltooligosaccharides (MOS) (Seung et al., 2017), but have no known enzymatic domains. However, they interact and act together with enzymes. For example, in Arabidopsis leaves, PTST2 interacts with SS4, and both proteins promote granule initiation in chloroplasts (Roldán et al., 2007; Seung et al., 2017). The elongation of soluble MOS is an important step towards initiating new starch granules (Nakamura, 2015; Seung and Smith, 2019; Mérida and Fettke, 2021). The plastid contains multiple enzymes that can elongate MOS - including SS4 and other starch synthase isoforms, as well as the plastidial α-glucan phosphorylase (PHS1 or PHO1). PHS1 catalyses a reversible reaction, either degrading α-1,4-linked glucan chains via a phosphorolysis reaction that releases glucose-1-phosphate (G1P) or extending the glucan chain using G1P as a substrate.

The role of PHS1 in granule initiation, or more broadly in starch metabolism, is not clear. In Arabidopsis, the complete loss of PHS1 does not affect growth, starch turnover (Zeeman et al., 2004) or granule number per chloroplast (Malinova et al., 2014). However, double mutants defective in PHS1 and other components of maltose metabolism have severe defects in growth and strong reductions in granule number per chloroplast (Malinova et al., 2014; Malinova and Fettke, 2017). A role for PHS1 in granule initiation was also inferred from rice PHS1 knockout mutants, which produced grains ranging from normal to shrunken; where shrunken grains with dramatic decreases in starch content were more prevalent when plants were grown at low temperatures (Satoh et al., 2008). Overall, these findings suggest PHS1 could play a role in granule initiation, but in the species examined, its importance is conditional on the presence of other mutations or temperature. More recent evidence suggests that PHS1 deficiency in potato and rice results in strong alterations in MOS levels (Dong et al., 2023; Flores-Castellanos and Fettke, 2023).

In this study, we aimed to discover the mechanism of B-type granule initiation. We looked for wheat endosperm proteins that interact with BGC1, and we discovered that it interacts with PHS1. We therefore investigated the role of PHS1 in the wheat endosperm, and we provide strong genetic evidence that PHS1 is required for normal initiation of B-type granules during grain development, but not for normal A-type granules. We therefore propose a model where A-and B-type granules initiate via distinct biochemical mechanisms, the latter requiring PHS1.

## Results

### BGC1 interacts with proteins involved in starch synthesis, including PHS1

To discover proteins involved in B-type granule initiation, we identified proteins that associate with BGC1 using affinity pulldown and mass spectrometry. We produced extracts from developing durum wheat endosperm (cultivar Kronos) harvested at 18 days post anthesis (dpa), a timepoint when B-type granule formation is actively occurring. Following coincubation with recombinant His-tagged BGC1, we pulled down BGC1 and associated proteins using anti-His beads. Mass spectrometry was used to identify interactors. We set a threshold for at least two-fold significant enrichment (*p* < 0.05) in the pulldown vs. controls (extracts incubated without *Ta*BGC1). The 118 proteins that met this criterion are presented in Supplemental Data 1.

Given the large number of proteins identified in the pulldown, we used the dataset to specifically assess which known proteins of starch metabolism pulled down with BGC1. There were 13 starch-related proteins in the filtered list (Table 1). We observed that MAR-binding filament-like protein (MFP1), a known interaction partner of PTST2 (the BGC1 ortholog) in Arabidopsis (Seung et al., 2018), was enriched with the highest abundance in the pulldown - providing strong indication that the pulldown was effective. MFP1 has been duplicated in cereals (Seung et al., 2018), and interestingly both paralogs of MFP1 in wheat (MFP1.1 and MFP1.2) were identified in the pulldown, suggesting both are present in the endosperm and can associate with BGC1. In addition, we found multiple starch branching enzyme 1 isoforms (SBE1). Only two proteins with glucan-elongating activity were detected in the pulldown: These were the plastidial α-glucan phosphorylase (PHS1) and starch synthase 2a (SS2a). In this study, we focused on the interaction between PHS1 and BGC1, given the prior knowledge that PHS1 elongates MOS and can conditionally act in granule initiation in other species (Satoh et al., 2008; Malinova et al., 2014), and because SS2a already has established roles in other aspects of starch synthesis, particularly in determining amylopectin structure (Morell et al., 2003). Notably, peptides matching SS4 were not detected in this experiment.

**Table 1:** Starch metabolic proteins significantly enriched in the BGC1 pulldown.

| Accession | Protein | Loc. | Abundance Ratio (pulldown/control) | Abundance Ratio P-Value |
| --- | --- | --- | --- | --- |
| TRITD3Av1G038460.3 | MAR binding filament-like protein 1.2 | C | 100 | 1E-17 |
| TRITD1Av1G054690.1 | MAR binding filament-like protein 1.1 | C | 100 | 1E-17 |
| TRITD3Bv1G047250.3 | MAR binding filament-like protein 1.2 | C | 93.506 | 1E-17 |
| TRITD1Bv1G062760.3 | MAR binding filament-like protein 1.1 | C | 56.176 | 1E-17 |
| TRITD7Bv1G229280.2 | Starch branching enzyme 1.3 | C | 45.482 | 1E-17 |
| TRITD6Bv1G086030.1 | Brittle1 transporter 1 | C | 26.309 | 1E-17 |
| TRITD7Av1G275430.16 | Starch branching enzyme 1.1 | C | 11.778 | 1E-17 |
| TRITD2Bv1G044170.1 | Disproportionating enzyme 2 | O | 3.363 | 0.0064 |
| TRITD7Bv1G229250.2 | Starch branching enzyme 1.1 | C | 2.949 | 0.0033 |
| TRITD5Bv1G140970.6 | Phosphoglucose isomerase | C | 2.563 | 0.0291 |
| **TRITD5Bv1G201740.3** | **Plastidial α-glucan phosphorylase** | **C** | **2.538** | **0.0211** |
| TRITD2Av1G048150.2 | Disproportionating enzyme 1 | C | 2.359 | 0.0428 |
| TRITD7Bv1G038900.1 | Starch synthase 2a | C | 2.308 | 0.0006 |

The predicted localisation (Loc.) of each protein as predicted by TargetP 2.0, as in the chloroplast (C) or other (O), is shown. The abundance ratio was calculated from the average abundances in three replicate pulldown and control reactions. These three replicates were used to calculate the P-value. The full dataset, including raw quantification values, are in Supplemental Data 1.

To confirm the interaction between BGC1 and PHS1 observed in wheat extracts, we carried out a pairwise immunoprecipitation assay using transiently expressed proteins in *Nicotiana benthamiana* leaves. We first confirmed that both PHS1 and BGC1 located to the chloroplast in this heterologous system (Figure 1A). PHS1 was located in the stroma, whereas BGC1 was located around starch granules, and in punctate structures previously described for the Arabidopsis ortholog (Seung et al., 2017; Seung et al., 2018). We then used YFP/RFP-tagged proteins for the immunoprecipitation. BGC1:RFP co-purified with PHS1:YFP in the IP with anti-YFP beads when the two fusion proteins were co-expressed (Figure 1B). BGC1:RFP did not purify in control reactions when co-expressed with a chloroplast-targeted YFP, nor did a chloroplast-targeted RFP co-purify with PHS1:YFP. This suggests that BGC1 specifically interacts with PHS1.

**Figure 1.**
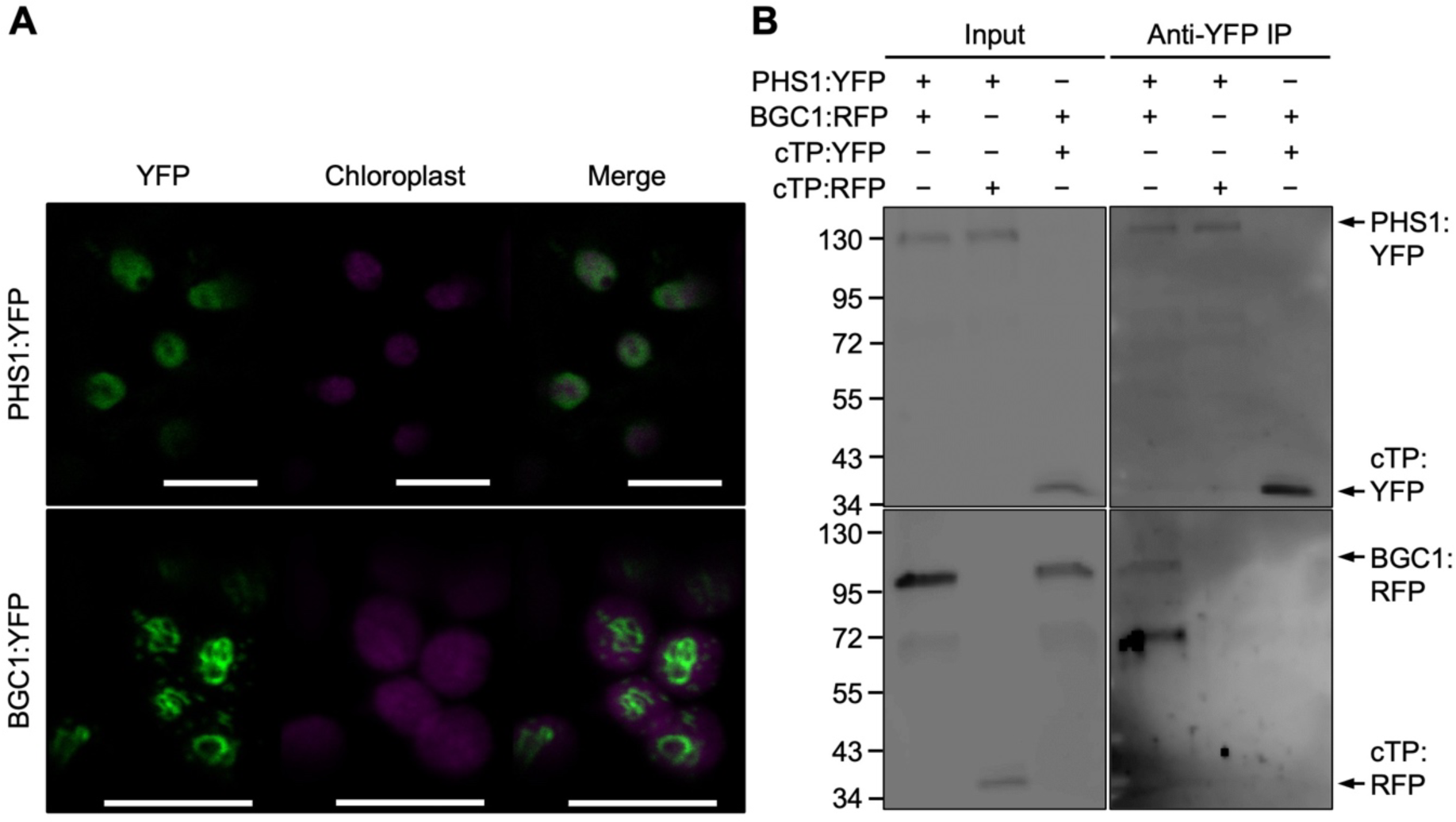
Interaction between wheat PHS1 and BGC1 in *Nicotiana benthamiana*. **A)** Plastidial localisation of YFP-tagged PHS1 and BGC1 transiently expressed in *Nicotiana* leaves. Bar = 10 µm. **B)** Pairwise immunoprecipitation (IP) of PHS1:YFP and BGC1:RFP co-expressed in *N. benthamiana* leaves, using anti-YFP beads. Input and IP samples were blotted with YFP (top panels) and RFP antibodies (bottom panels). Chloroplast-targeted YFP (cTP:YFP) and RFP (cTP:RFP) were used as controls to exclude unspecific binding to the fluorescent protein tags.

### PHS1 is encoded on group 5 chromosomes in wheat

To study the role of PHS1 in wheat, we first identified the gene models corresponding to PHS1. BLAST searches using the Arabidopsis PHS1 on Ensembl plants (Kersey et al., 2018) against the bread wheat (*Triticum aestivum*) reference genome (cultivar Chinese Spring, RefSeq v1.1) (Appels et al., 2018) showed three homeologs of PHS1 encoded on group 5 chromosomes. These were *PHS1-A1* (TraesCS5A02G395200), *PHS1-B1* (TraesCS5B02G400000) and *PHS1-D1* (TraesCS5D02G404500) (Figure 2A). In the durum wheat (*Triticum turgidum ssp. durum*) reference genome (cultivar Svevo, Svevo.v1)(Maccaferri et al., 2019), these corresponded to *PHS1-A1* (TRITD5Av1G205670) and *PHS1-B1* (TRITD5Bv1G201740). The primary transcripts at all these loci had 15 exons and 14 introns; with the exception of *PHS1-B1* in durum wheat which due to poor gene model prediction, we reannotated based on the gene model in Chinese Spring. After the reannotation, the predicted amino acid sequences for PHS1 between bread and durum wheats were 99.9% and 99.0% identical for *PHS1-A1* and *PHS1-B1*, respectively.

**Figure 2.**
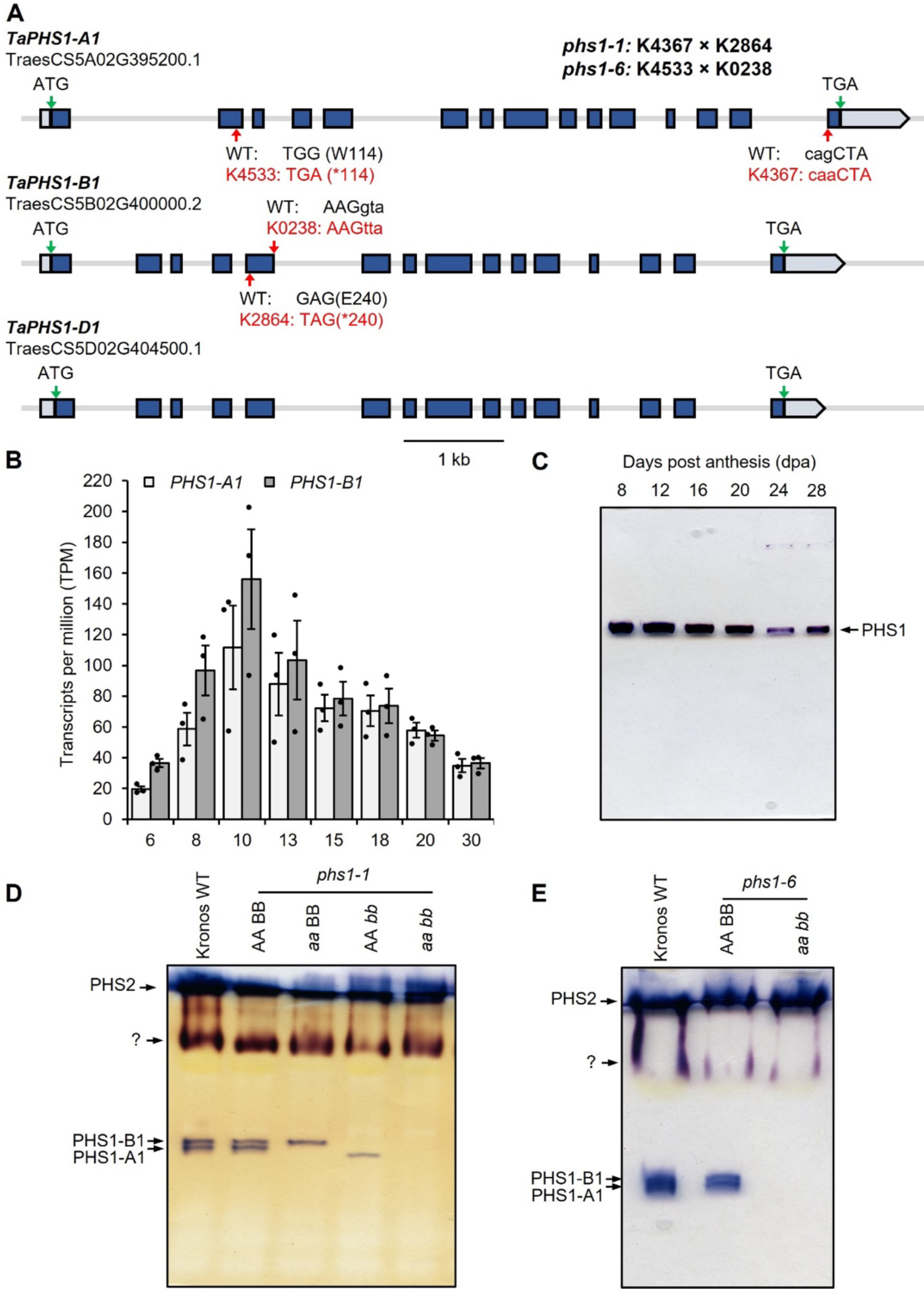
Expression of wheat *PHS1* loci and generation of loss-of-function mutants. **A)** Gene models of *PHS1* homoeologs in Chinese Spring. Exons are depicted in dark blue and the 5’ and 3’ UTRs are depicted in light blue. The position of the start (ATG) and stop (TGA) codons are indicated with green arrows. The position of the mutations in the TILLING mutants are indicated with red arrows. **B)** Expression levels of *PHS1* homeologs in the endosperm of durum wheat across different stages of grain development. Data are from the RNAseq study by Chen et al. (2022). Values are in transcripts per million (TPM) and are means ± SEM from n = 3 replicates per time point. **C)** Native PAGE of phosphorylase activity. Crude extracts were produced from dissected endosperms across different stages of grain development and were separated on 7.5% polyacrylamide gels containing 0.3% glycogen (∼180 µg protein per well). Activity bands were visualised after incubation with G1P by staining in Lugol’s iodine solution. **D)** Native PAGE of phosphorylase activity using extracts of leaves from 4-week-old seedlings of the wild type (WT), *phs1-1* single (*aa* BB and AA *bb*) and double (*aa bb*) mutants, and wild-type segregant controls (AA BB). The gel was run as for C) **E)** As for **D)**, but with *phs1-6* mutants.

We then compared the expression patterns of PHS1 and BGC1 during grain development. We used our previously generated RNAseq dataset for endosperm development in durum wheat cultivar, Kronos (Chen et al., 2022). PHS1 expression was detectable at all stages of grain development tested between 6-30 dpa but was particularly strong between 8-13 dpa (Figure 2B). No differences in expression level or pattern were observed between the A- and B-homeologs. PHS1 activity levels in endosperm extracts were visualised on native PAGE gels. Consistent with the transcript levels, robust PHS1 activity was detected at all timepoints tested between 8-28 dpa, but was stronger at the earlier than later timepoints, especially at 12 dpa (Figure 2C). To assess PHS1 expression in different tissues, we used public data for bread wheat cultivar Chinese Spring in the wheat expression browser (Borrill et al., 2016). *PHS1* transcripts were detectable in leaves/shoots, roots, spike, and grains, and all homoeologs showed similar levels of expression across all examined tissues (Supplemental Figure 1).

### The phs1 mutant of wheat has no obvious effects on vegetative growth

To study the function of PHS1, we used the wheat in silico TILLING resource (http://www.wheat-tilling.com)(Krasileva et al., 2017) to identify mutants in tetraploid durum wheat (*T. turgidum* cv. Kronos) with mutations in either homoeolog. The Kronos4533 (K4533) and Kronos4367 (K4367) lines had premature stop or splice acceptor mutations respectively in *PHS1-A1*, whereas Kronos0238 (K0238) and Kronos2864 (K2864) had premature stop and splice donor mutations for *PHS1-B1* (Figure 2A). To combine A- and B-homoeolog mutant alleles, K4367 was crossed with K2864 to generate *phs1-1* mutant lines, and K4533 was crossed with K0238 generate *phs1-6* mutant lines. The AA BB (wild-type sibling control), *aa* BB and AA *bb* (single mutants), and *aa bb* (double mutant) genotypes were selected in the F2 generation using KASP genotyping.

We verified the effect of the mutations on PHS1 activity using native PAGE gels, using proteins extracted from leaf tissue. Unlike in the endosperm (Figure 2C), we observed four different phosphorylase activities in leaf protein extracts (Figure 2D and E). Further, due to the lower activity in leaves compared to endosperm, we could distinguish two closely migrating PHS1 bands for leaf extracts. Both bands were absent in the *phs1-1* and *phs1-6 aa bb* double mutants, and the two *aa* BB and AA *bb* single mutants of *phs1-1* were missing either one of the two bands. We concluded that the two bands represent PHS1-A1 and PHS1-B1 activities, and the selected mutations for *phs1-1* and *phs1-6* mutant lines eliminate all detectable PHS1 activity. The strong upper activity bands on the gels likely correspond to the cytosolic phosphorylase, PHS2 (Schupp and Ziegler, 2004).

Overall, we observed no detectable effect of the *phs1* mutants on plant growth in glasshouses (Figure 3A). However, in some experiments, plants from all *phs1-1* lines (including the mutants and the wild-type siblings) were slightly shorter than wild type plants. This is likely due to background mutations rather than the loss of PHS1 activity, since it was also observed in the wild-type sibling from the *phs1-1* cross but not in any of the *phs1-6* lines. Similarly, there was no effect of *phs1* mutations on the grain yield per plant, but a decrease was seen in all *phs1-1* lines (Figure 3B). Again, there was no significant difference in grain yield per plant or seed number between the *phs1-1* double mutant and its wild-type sibling control, or between the *phs1-6* double mutant and its wild-type sibling control (Figure 3B). All lines produced seeds that appeared normal (Figure 3C), and we did not observe any significant effects of *phs1* mutations on thousand grain weight or grain size (Figure 3D, E). Since the rice mutant is reported to have a significant proportion of shrunken seeds (Satoh et al., 2008; Dong et al., 2023), we also assessed the distribution of grain sizes harvested from the mutants, but the proportion of small seeds was not greater in the mutants compared to the WT (Supplemental Figure 2). We therefore conclude that loss of PHS1 activity does not affect vegetative growth, seed yield or seed morphology under our conditions.

**Figure 3.**
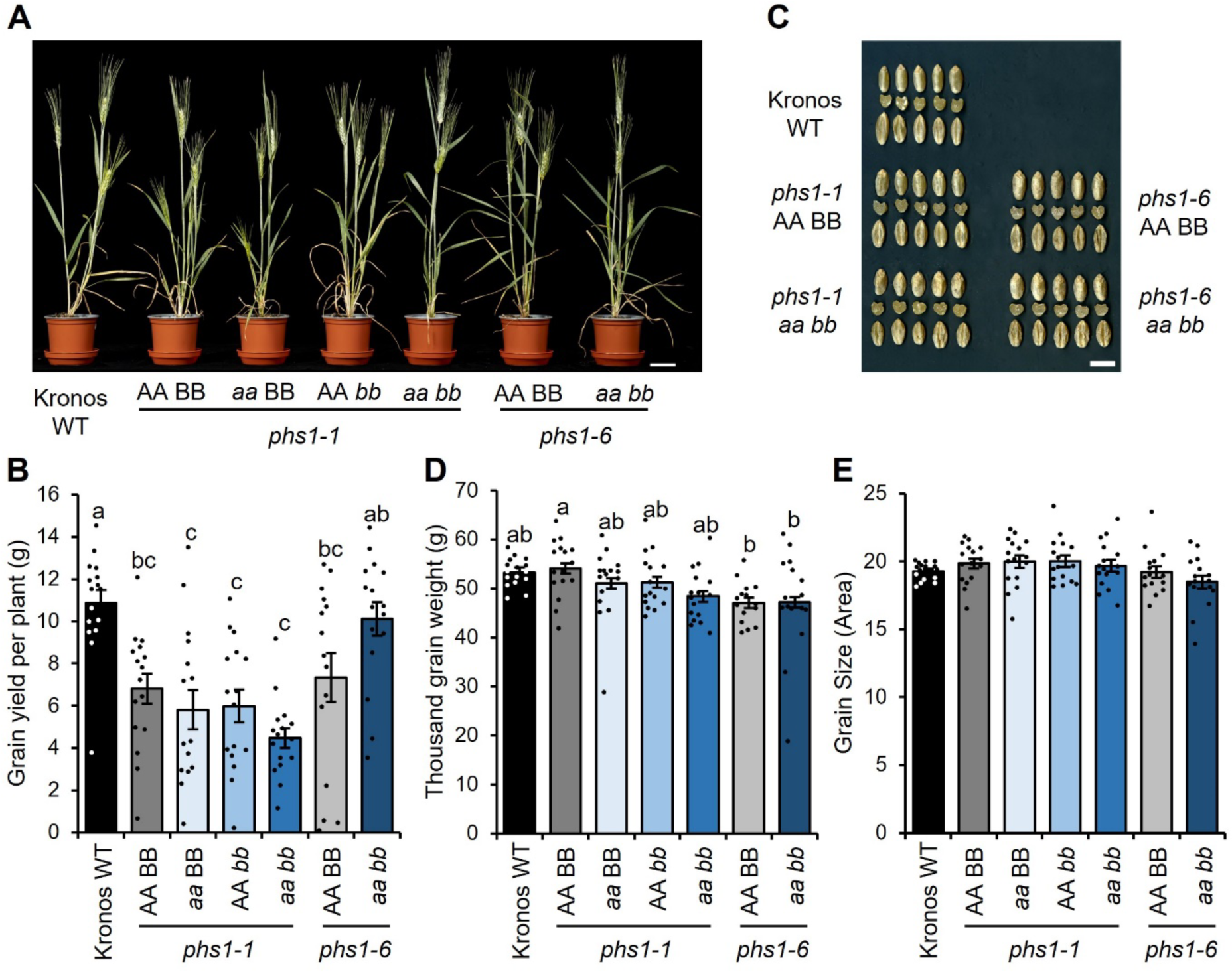
Plant and grain phenotypes of *phs1* mutants. The wild-type sibling control (AA BB), single mutants (*aa* BB and AA *bb*) and the double mutant (*aa bb*) from *phs1-1* and *phs1-6* lines were compared with the Kronos wild type (WT). **A)** Photograph of plants at maturity. Bar = 5 cm. **B)** Grain yield per plant. **C)** Photographs of grains showing both dorsal and ventral sides, as well as a cut section through the middle of the grain. Bar = 1 cm. **D)** Thousand grain weight. **E)** Grain size measured as 2-D area. For panels B, D and E, data represent the mean *±* SEM from *n* = 15-16 plants. Values with different letters are significantly different under a one-way ANOVA and Tukey’s post hoc test at *p* < 0.05. There were no significant differences between genotypes in panel E.

### PHS1 is required for normal B-type granule size and number

We investigated the effect of *phs1* mutations on starch synthesis in the endosperm. First, we measured the total starch content of grains. The *phs1* double mutants had identical starch content to the wild-type segregant control and the wild type (Table 2). Interestingly, starch in both the *phs1-1* and *phs1-6* double mutants had a small but significant reduction in amylose content, suggesting that PHS1 is required for normal amylose content in wheat. The reduced amylose content was not observed in the *phs1-1* single homeolog (*aa* BB, AA *bb*) mutants.

**Table 2.** Starch and amylose content of *phs1* mutants.

| **Table 2 - Starch and amylose content of *phs1* mutants** |  |  |
| --- | --- | --- |
|  | Starch content (%) | Amylose content (%) |
| **Experiment 1 – *phs1-1* single and double mutants** |  |  |
| Kronos WT | 47 ± 2<sup>ab</sup> | 28.6 ± 2.4<sup>a</sup> |
| *phs1-1* AA BB | 44 ± 2<sup>ab</sup> | 28.8 ± 0.8<sup>a</sup> |
| *phs1-1* *aa* BB | 39 ± 3<sup>b</sup> | 26.7 ± 0.6<sup>a</sup> |
| *phs1-1* AA *bb* | 40 ± 1<sup>b</sup> | 26.6 ± 1.3<sup>a</sup> |
| *phs1-1* *aa* *bb* | 49 ± 2<sup>a</sup> | 21.1 ± 0.6<sup>b</sup> |
| **Experiment 2 – *phs1-1* and *phs1-6* double mutants and *bgs1-1*** |  |  |
| Kronos WT | 55 ± 2<sup>a</sup> | 21.5 ± 0.3<sup>d</sup> |
| *phs1-1* AA BB | 49 ± 2<sup>a</sup> | 20.8 ± 0.2<sup>cd</sup> |
| *phs1-1* *aa* *bb* | 44 ± 1<sup>a</sup> | 17.8 ± 0.2<sup>ab</sup> |
| *phs1-6* AA BB | 54 ± 1<sup>a</sup> | 19.3 ± 0.2<sup>bc</sup> |
| *phs1-6* *aa* *bb* | 44 ± 3<sup>a</sup> | 17.9 ± 0.2<sup>ab</sup> |
| *bgs1-1* | 43 ± 4<sup>a</sup> | 16.9 ± 1<sup>a</sup> |
| Starch content was quantified relative to the dry weight of whole flour. Values are the mean ± SEM on grains from *n* = 4 different plants. Amylose content was measured on purified starch. Values are the mean ± SEM on starch extracted from *n* = 4-6 different plants. Values with different letters are significantly different under a one-way ANOVA and Tukey's posthoc test at *p* < 0.05. Note that the plants for the two experiments were grown independently, and therefore the statistical analyses compare genotypes within each experiment. |  |  |

We then purified starch granules from mature grains and examined their morphology using Scanning Electron Microscopy (SEM). Interestingly, starch from the *phs1* double mutants, *phs1-1* and *phs1-6* had visibly fewer B-type granules than the wild type (Figure 4A and Supplemental Figure 3A). Quantification of granule size distribution on the Coulter counter confirmed that the granule size distribution (plotting relative volume over diameter) was drastically altered in both *phs1-1* and *phs1-6* double mutants, with the B-type granule peak being smaller compared to the WT controls, and shifted towards the larger sizes (Figure 4B and Supplemental Figure 3B). Using curve-fitting on these plots, we calculated that both *phs1-1* and *phs1-6* double mutants had less than half of the B-type granule content (by volume) compared to the wild-type and single mutants, despite having significantly larger B-type granules (Figure 4C, D and Supplemental Figure 3C, D). As the large size of the B-type granules in the double mutant affects the area of the B-type granule peak when plotted on a volume basis, we also plotted the size distributions on a relative number basis for the *phs1-1* double mutant (Figure 4E), which showed a large shift in the proportion of A- vs. B-type granules in the double mutant. The granule size distributions in the *phs1-1* single homeolog mutants were identical to the WT controls, suggesting that mutations in both PHS1 homeologs are necessary to alter this phenotype (Figure 4A-E).

**Figure 4.**
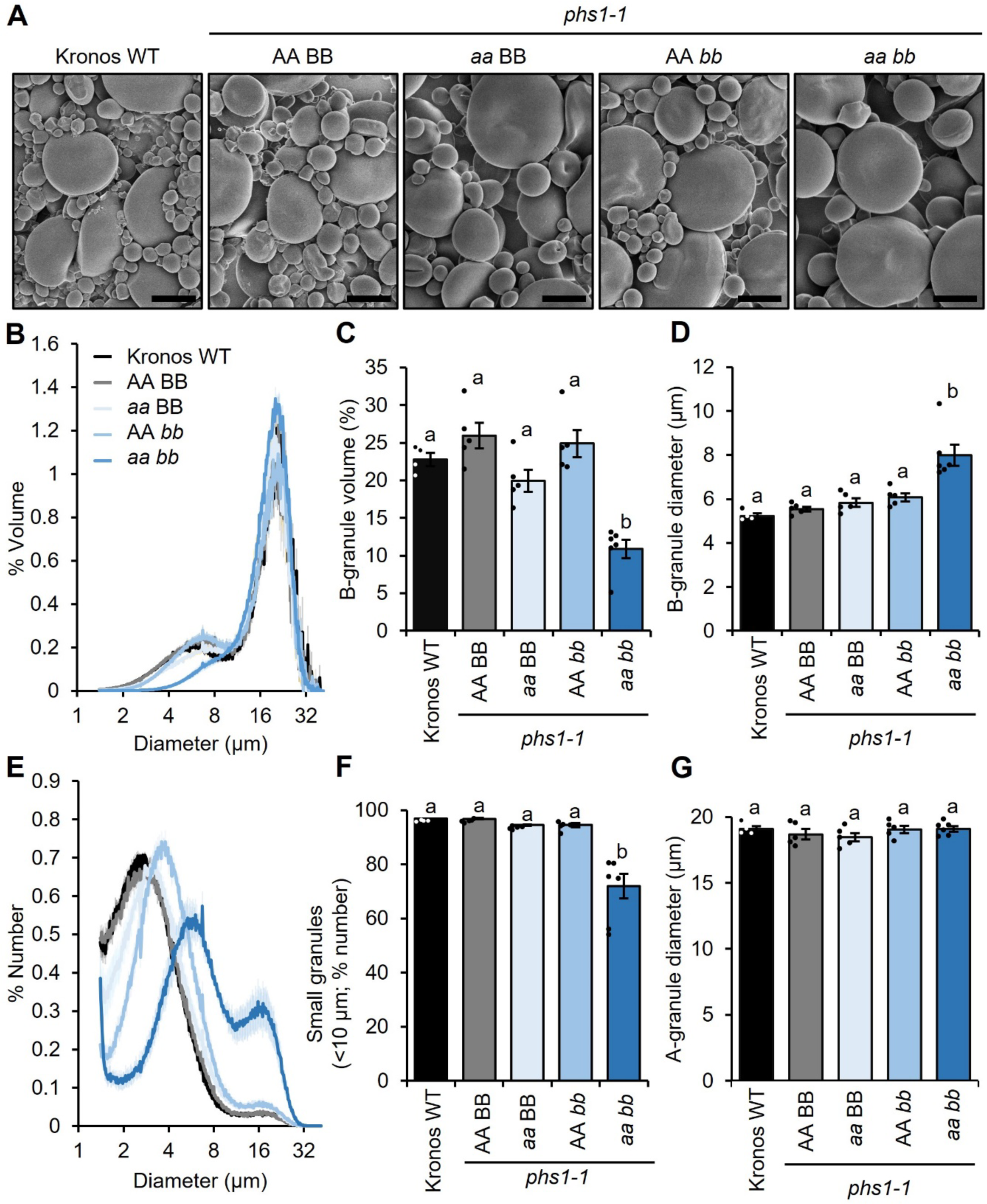
Endosperm starch from the *phs1-1* mutant has fewer and larger B-type granules. **A)** Scanning electron micrographs of purified endosperm starch. Bar = 10 µm. **B)** Granule size distributions were determined using a Coulter counter, and the data were expressed as relative % volume (of total starch) vs. granule diameter plots. **C** and **D)** B-type granule volume (% of total starch) and the average diameter of B-type granules were extracted from the relative volume vs. diameter plots by fitting a bimodal mixed normal distribution. **E)** Same as B, but expressed as relative starch granule number (% of total granule number) vs. diameter plots. **F)** The percentage of small granules by number (smaller than 10 µm) were calculated from the Coulter counter data. **G)** The average diameter of A-type granules was calculated from the relative volume vs. diameter plots, as for panels C and D. For panels B-G, plots show the mean from the analysis of n=4-6 replicate starch extractions, each from grains from a separate plant. The shading (on panels B and E) and error bars (on panels C, D, F, G) represent the SEM. Values with different letters are significantly different under a one-way ANOVA with Tukey’s post-hoc test (p < 0.05).

As the large size of the B-type granules affects the calculation of B-granule content on a volumetric basis, we also investigated whether the number of B-type granules were affected in the double mutant. Curves cannot be reliably fitted to number-diameter plots from the Coulter counter due to the very small size of the A-type granule peak. So instead, as done previously (Chia et al., 2020), we calculated the percentage of starch granules that were smaller than <10 µm, which includes B-type granules and some smaller A-type granules. There was a significant decrease in the relative number of small granules in the double mutant compared to the controls (Figure 4F and Supplemental Figure 3E). By contrast to B-type granules, the size of A-type granules was unaffected in both *phs1-1* and *phs1-6* double mutants (Figure 4G and Supplemental Figure 3F).

Taken together, these data suggest that loss of PHS1 results in the synthesis of fewer, but larger, B-type granules. This, together with its interaction with BGC1, points to a role for PHS1 in the initiation of B-type granules which we investigate further below. Additionally, PHS1 is required for normal amylose content in wheat grains.

### PHS1 acts during B-type granule initiation in grain development

To examine if the loss of PHS1 specifically affected B-type granule formation during grain development, we measured starch content, granule number and granule size distributions in developing endosperm. We dissected endosperms from developing grains of the *phs1-1* and *phs1-6* double mutants, as well as their corresponding wild-type controls. Similar to in mature grains, we did not observe significant differences in total starch content between the mutants and wild-type controls at any time point during grain development (Figure 5A). We then used the Coulter counter to quantify the number of starch granules per mg of endosperm, as well as to assess granule size distributions. The number of starch granules did not differ between the mutants and the wild type at 8 or 14 dpa. However, at 18 and 22 dpa, both the *phs1* mutants had significantly fewer starch granules than the wild-type controls (Figure 5B).

**Figure 5.**
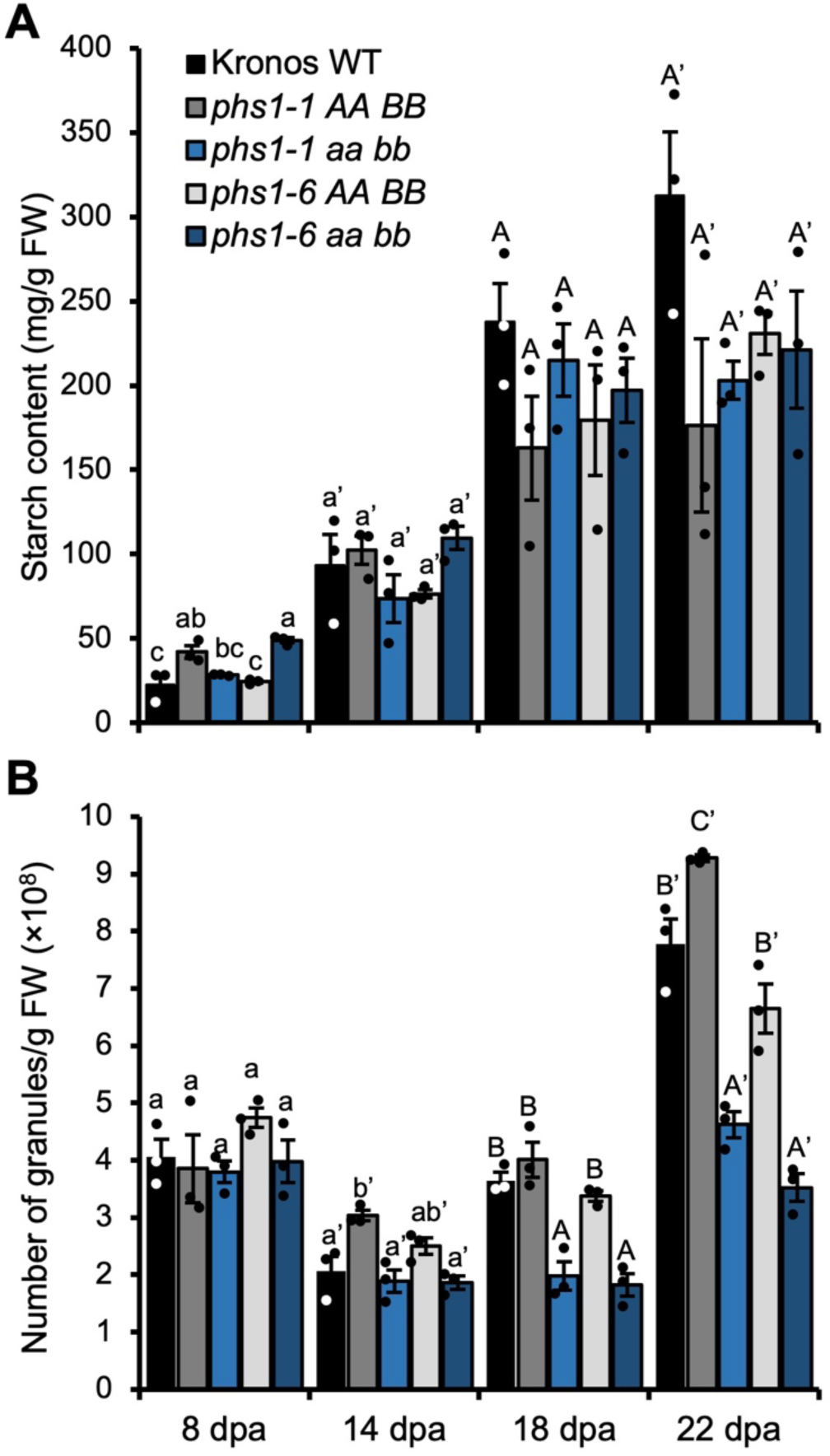
Loss of PHS1 affects granule number in mid-grain development but not total starch content. The endosperm was dissected from developing grains of WT, *phs1-1* and *phs1-6* double mutants and corresponding wild-type controls, harvested at 8-, 14-, 18- and 22-days post anthesis (dpa), with n = 3 individual plants for each genotype per time point **A)** Starch content of the endosperm. Values are expressed relative to the fresh weight of the dissected endosperm. **B)** Starch granule number in the endosperm. Starch was purified from dissected endosperm and the number of granules was determined using a Coulter counter running in volumetric mode (analysing 2 mL of the suspension). Values are expressed relative to the fresh weight of the dissected endosperm. Values with different letters are significantly different under a one-way ANOVA with Tukey’s post-hoc test (p < 0.05).

Since B-type granules initiate typically between 15-20 dpa, and only A-type granules are present before this timepoint, it is likely that the *phs1* mutants contain normal numbers of A-type granules (Figure 6A, B), but have fewer B-type granules than the wild type (Figure 6C, D). This hypothesis was also strongly supported by the granule size distributions from the Coulter counter and in SEM (Figure 6A-H).

**Figure 6.**
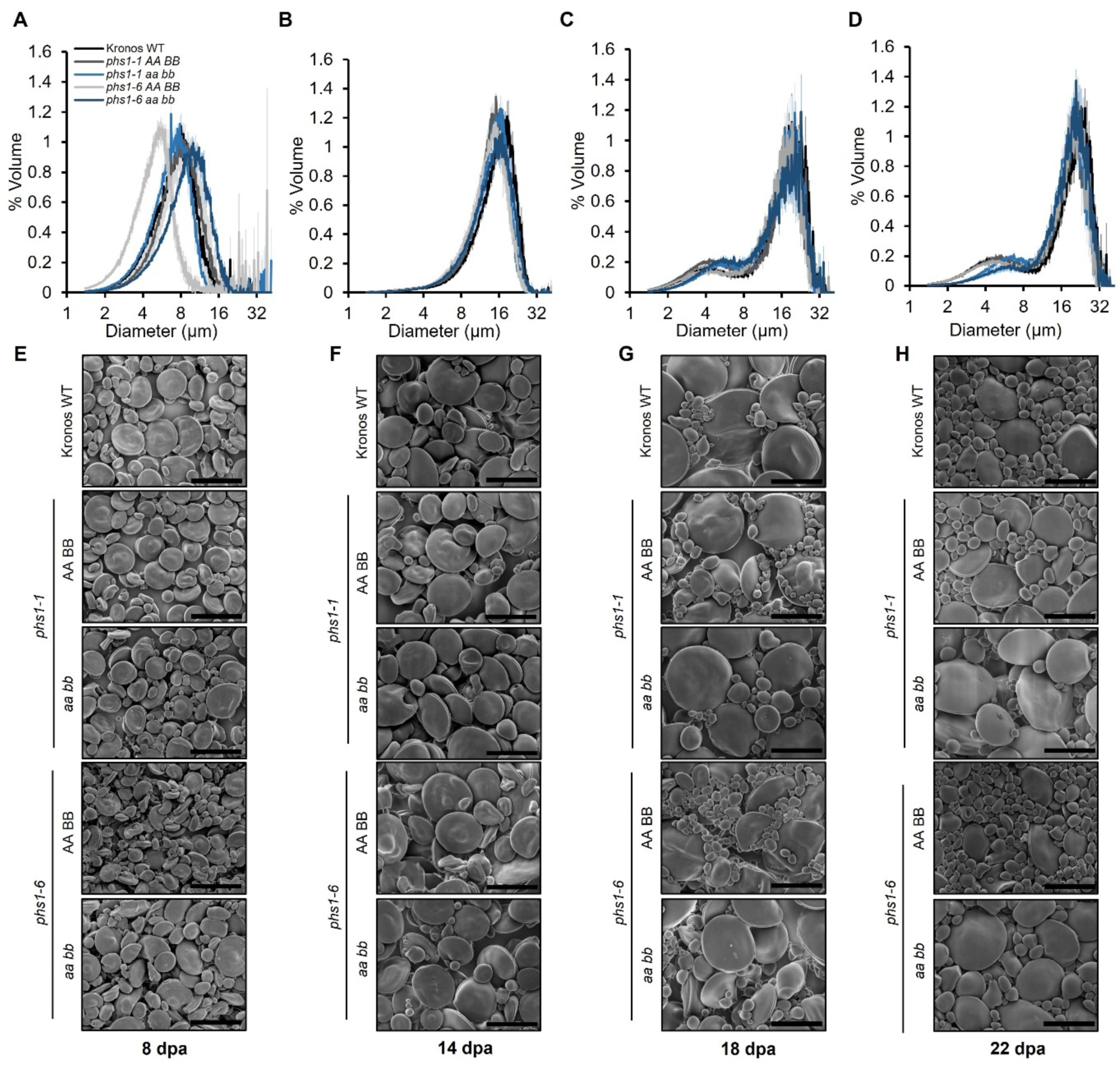
PHS1 affects granule size distributions during mid-grain development. The endosperm was dissected from developing grains of WT, *phs1-1* and *phs1-6* double mutants and corresponding wild-type controls, harvested at **A-E**) 8 days post anthesis (dpa), **B-F**) 14 dpa, **C-G**)18 dpa and **D-H**) 22 dpa, with n = 3 individual plants for each genotype per time point **A-D)** Granule size distributions were analysed on the Coulter counter and the data were expressed as relative % volume (of total starch) vs. granule diameter plots. The shading represents the ± SEM. **E-H)** Starch granule morphology observed using scanning electron microscopy. Bars = 15 µm.

The distributions were unimodal in the mutants and the wild-type controls at the 8 and 14 dpa timepoints (Figure 6A, B, E, F). However, at the 18 and 22 dpa timepoints, the distributions turned bimodal – suggesting all genotypes could initiate B-type granules (Figure 6C, D, G, H). However, the B-type granule peaks were smaller and shifted towards the larger sizes in the *phs1* mutants (Figure 6C, D). Larger, and fewer B-type granules were observed in the SEM images of starch from these timepoints (Figure 6G, H). There were no defects in A-type granule morphology at any timepoint. These data suggest that PHS1 is required for normal B-type granule initiation, but not for the correct number and morphology of A-type granules (Figure 6).

Our finding that PHS1 is required solely for B-type granule initiation was further supported in a genetic approach. Reduced gene dosage of BGC1 affects the number of B-type granules – such as in the *bgc1-1* mutant of durum wheat, Kronos, which almost has no B-type granules due to a loss-of-function mutation in the *BGC1-A1* homeolog and a missense mutation in the *BGC1-B1* homeolog (Chia et al., 2020). We crossed this *bgc1-1* line to the *phs1-1* line, and isolated a *bgc1-1 phs1-1* quadruple mutant in the F2 generation. We also isolated from the cross a wild-type sibling control, and *bgc1-1* and *phs1-1* mutant siblings as controls. We purified starch granules from mature grains of these lines and examined their morphology using Scanning Electron Microscopy (SEM). Starch from the *bgc1-1* and *bgc1-1 phs1-1* mutants had visibly fewer B-type granules than the wild type, wild-type sibling control, and *phs1-1* mutant siblings (Figure 7A). Coulter counter analysis of granule size distributions showed that the reduction of B-type granule number was stronger in *bgc1-1* than in *phs1-1* (Figure 7B), since *bgc1-1* had almost no detectable B-type granule peak (when plotted by volume), whereas *phs1-1* had a B-type granule peak that was between that of the wild-type controls and *bgc1-1*. Importantly, the granule size distribution of *bgc1-1 phs1-1* quadruple mutants was almost identical to *bgc1-1*. This suggests that the loss of PHS1 in the *bgc1-1* background (which only has A-type granules), has no further effect on starch granule size distributions. Using curve-fitting on these plots, we calculated that the size of A-type granules was unaffected by the *phs1* mutations in the *bgc1-1* background (Figure 7C). Next, we calculated the percentage of starch granules that were smaller than <10 µm. There was a significant decrease in the relative number of small granules in the *bgc1-1 phs1-1* quadruple mutant and *bgc1-1* and *phs1-1* mutant siblings compared to the wild-type controls (Figure 7D). Interestingly, like the *phs1* double mutants, we observed that *bgc1-1* also has a small reduction in amylose content (Table 2). Taken together, our results suggest PHS1 has an exclusive role in B-type granule initiation, and does not appear to be required for A-type granule initiation.

**Figure 7.**
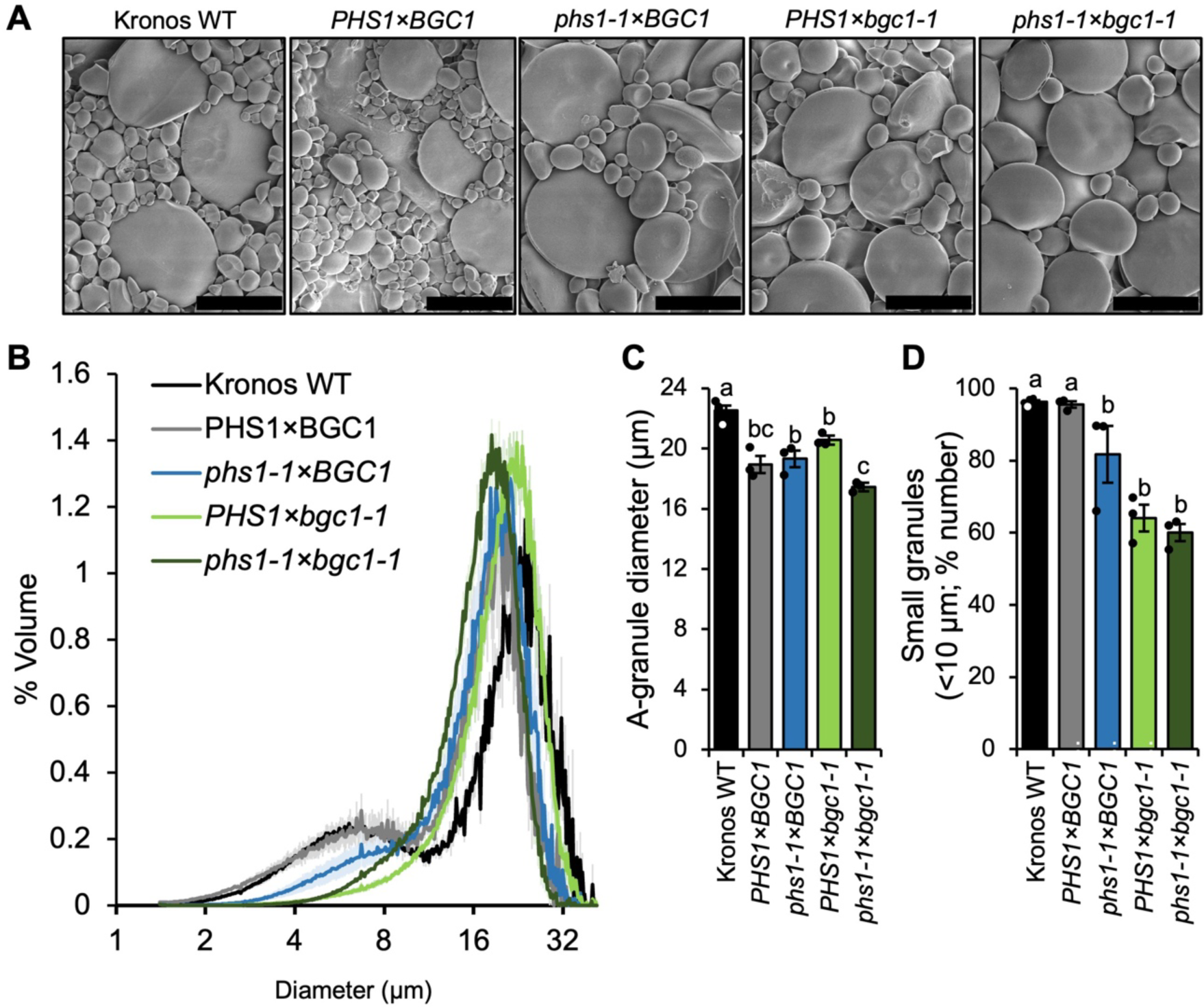
Loss of PHS1 does not affect granule size distribution in the *bgc1-1* mutant. **A)** Scanning electron micrographs of purified endosperm starch. Bar = 15 µm. **B)** Granule size distributions were determined using a Coulter counter, and the data were expressed as relative % volume (of total starch) vs. granule diameter plots. **C)** A-type granule diameter extracted from the relative volume vs. diameter plots by fitting a bimodal mixed normal distribution, except for genotypes with *bgc1-1*, where a unimodal distribution was fitted. **D)** The percentage of small granules by number (smaller than 10 µm) were calculated from the Coulter counter data. For panels B-D, plots show the mean from the analysis of n=3 replicate starch extractions, each from grains from a separate plant. The shading (in panel B) and error bars (in panels C and D) represent the ±SEM. Values with different letters are significantly different under a one-way ANOVA with Tukey’s post-hoc test (p < 0.05).

### Loss of PHS1 in wheat does affect total MOS levels in the endosperm

Given recent evidence that PHS1 deficiency in potato and rice results in strong MOS accumulation in tubers and endosperm (Dong et al., 2023; Flores-Castellanos and Fettke, 2023), we investigated whether there was MOS accumulation in the developing endosperm of wheat *phs1* mutants. However, levels of MOS, soluble glucans and other sugars were unaltered in the *phs1* mutants: We quantified the total soluble glucans in perchloric acid extracts. For all genotypes, soluble glucans were highest at 8 dpa, and decreased in abundance during grain development, but were always relatively low compared to starch content (approx. 10% of starch content at 8 dpa, and <0.001% at 22 dpa) (Supplemental Figure 4). There were no consistent differences between the *phs1* mutants and their controls. We then quantified the methanol precipitable fraction of soluble glucans (phytoglycogen and long MOS), and calculated the non-precipitable fraction (short MOS) as the difference between the total soluble glucan and precipitable glucans. Like the total glucans, these decreased in abundance as the grain developed, and there were no consistent differences between the *phs1* mutants and controls (Supplemental Figure 4). Additional High Performance Anion Exchange Chromatography with Pulsed Amperometric Detection (HPAEC-PAD) analyses largely confirmed this result, since maltose and other detectable MOS did not change in abundance or chain length distribution pattern in the *phs1* mutants (Supplemental Figure 5). In addition, glucose, fructose and sucrose levels remained unchanged in the mutants at all timepoints (Supplemental Figure 5).

### Loss of PHS1 does not affect starch granule number or starch turnover or in leaves

To test whether the loss of PHS1 affected leaf starch metabolism, we first quantified the total starch content in leaves of seedlings. There was no significant difference in starch content between the wild type and mutants, at either the end of day or end of night (Figure 8A; Supplemental Figure 6A). This indicates that PHS1 is not required for normal levels of starch accumulation and nocturnal starch turnover. To examine the number and morphology of starch granules in chloroplasts, we examined semi-thin sections of leaves using light microscopy (Figure 8B; Supplemental Figure 6B). There were no discernible differences in starch granule number per chloroplast in any of the genotypes. Like in Arabidopsis (Zeeman et al., 2004; Malinova et al., 2014), PHS1 appears to be dispensable for normal starch turnover in leaves.

**Figure 8.**
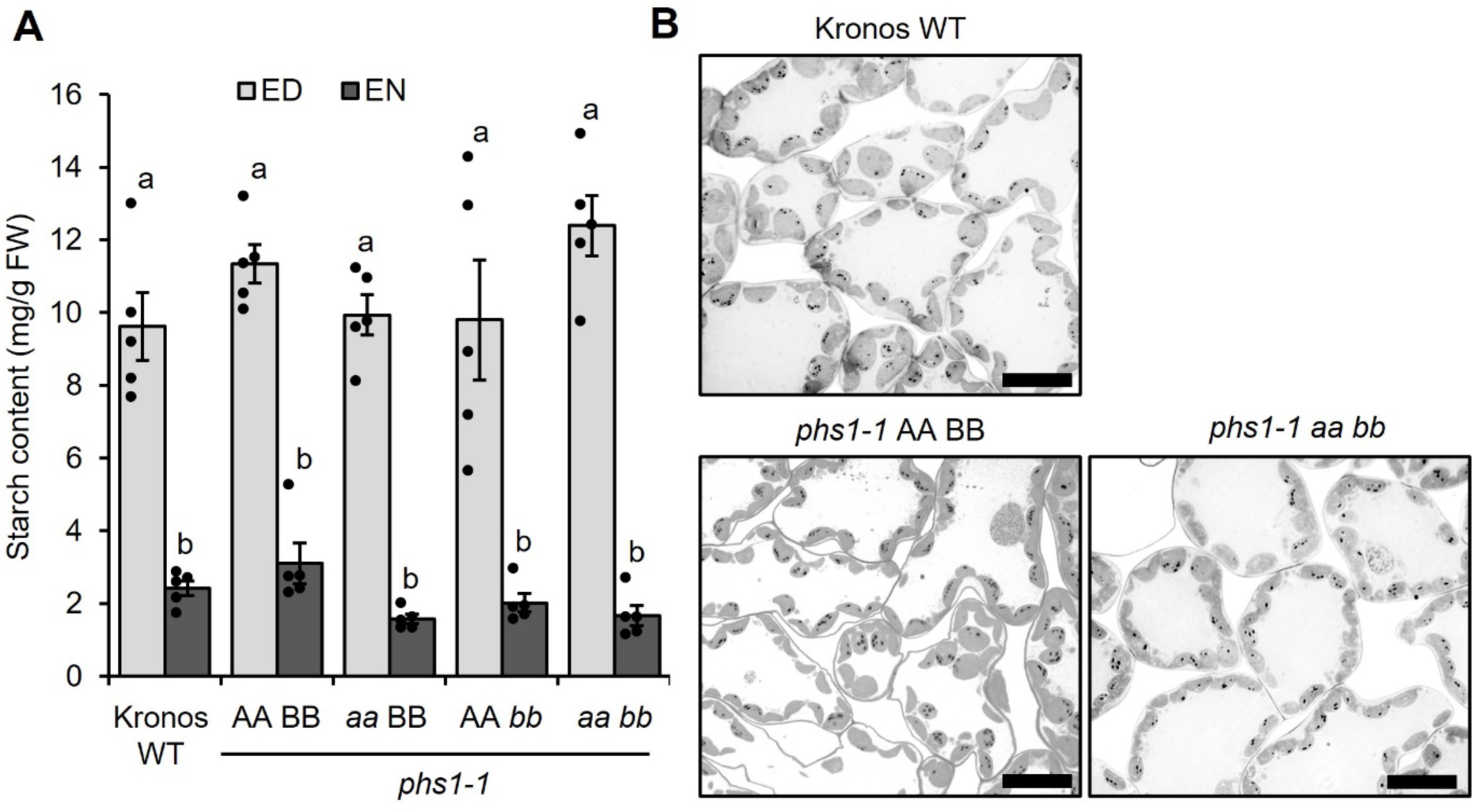
Leaf starch content of *phs1-1* mutants. The wild-type segregant control (AA BB), single mutants (*aa* BB and AA *bb*) and double mutants (*aa bb*) were compared with the Kronos wild type (WT). **A)** Total starch content. Seedlings were grown for two weeks under 16 h day/8 h night and harvested at the End of Day (ED) and End of Night (EN). Values are the mean±SEM from *n* = 5 plants (each represented by a data point), and those with different letters are significantly different under a one-way ANOVA and Tukey’s posthoc test at p < 0.05. **B)** Light micrographs of leaf sections stained with Periodic Acid-Schiff (PAS) staining to visualise starch granules. Leaf segments were harvested halfway along the length of the older blade in two-week-old seedlings were harvested at the ED, prior to fixation and embedding.

## Discussion

### Distinct mechanisms for A- and B-type granule initiation in wheat

After discovering PHS1 in screen for protein-protein interactions for BGC1 in wheat, we demonstrate that PHS1 is required for normal starch granule number and size distribution in the wheat endosperm, due to a specific role of the enzyme in B-type granule formation. We presented three lines of evidence to support this role: Firstly, mutants of durum wheat defective in *phs1* had fewer but larger B-type granules, with no differences in the number or size of A-type granules (Figure 4-5). Secondly, granule size distributions in the mutants were identical to those of the wild type during the early stages of grain development, but deviated from the wild type at 18 dpa - the timepoint after B-type granules started forming in the wild type (Figures 5-6). Finally, the *bgc1-1 phs1-1* double mutant had similar granule size distributions to the *bgc1-1* mutant, suggesting that the further loss of PHS1 in a background that only has A-type granules has no effect on granule size distribution (Figure 7). Thus, PHS1 appears to be only required for normal B-type granule formation in the wheat endosperm, and not for A-type granule formation.

Based on our findings, we present a model where A- and B-type granule initiation occur via distinct mechanisms involving different enzymes (Figure 9). B-type granule initiation is fundamentally different to A-type granule initiation in that B-type granules initiate within amyloplasts that already contain an A-type granule. They can therefore be considered “secondary” granule initiations. The A-type granule can act as a source of substrates that can prime B-type granule formation, particularly as MOS are released through the process of amylopectin trimming by Isoamylase (ISA) (Mouille et al., 1996; Tickle et al., 2009). By contrast, A-type granules are the first granules to form within each amyloplast during early grain development, and are more likely to require *de novo* primer formation. We propose that in wheat, PHS1 may only be important in the secondary B-type granule initiations, perhaps as it processes existing MOS substrates in the plastid in a manner that allows B-type granule initiation.

**Figure 9.**
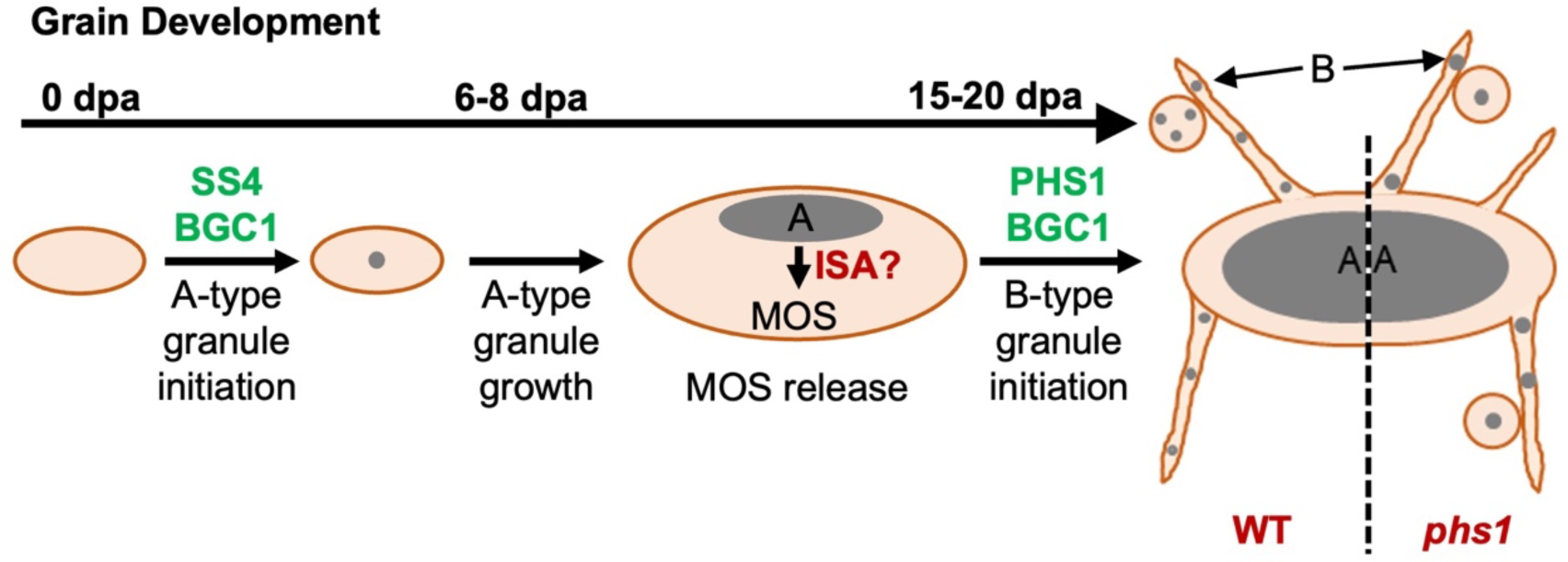
Model of PHS1 action in B-type starch granules initiation in wheat endosperm. During wheat grain development, a single A-type granule initiates in each amyloplast during early grain development by around 6-8 days post anthesis (dpa) and grows. B-type granules initiate later in grain development, at least partially in stromules and vesicle-like amyloplasts at around 15-20 dpa. We propose PHS1 acts on maltooligosaccharides (MOS), likely released from A-type granules by isoamylases (ISA), to initiate B-type granules. This process is disrupted in mutants lacking a functional PHS1 protein, resulting in fewer initiations of B-type starch granules. Fewer B-type granules means each one has a greater share of substrates for granule growth, leading to larger B-type granules in the mutant at grain maturity.

In this process, PHS1 may be assisted by BGC1. The number of B-type granules in wheat can be greatly reduced either through a loss of PHS1 function, or a reduction in BGC1 function (Figure 7)(Chia et al., 2020). BGC1 contains a glucan-binding CBM48 domain but no known enzymatic domain, and so is likely to act by influencing the substrate binding of its interacting enzymes, as previously proposed for PTST2 in Arabidopsis (Seung et al., 2017). Since BGC1 interacts with PHS1 in wheat, it could influence the substrates available for further elongation by PHS1 (Figure 1, Table 1). However, it appears that BGC1 is more strictly required for B-type granule formation than PHS1. B-type granules are almost absent in the *bgc1-1* mutant, whereas the *phs1* mutant can still initiate some B-type granules (Figure 7). It is very likely that this reflects the degree of redundancy in the function of both proteins. BGC1 is the only PTST2/3 ortholog in wheat and may be indispensable for B-type granule formation. PHS1 is the only plastidial phosphorylase, but it is not the only plastidial enzyme that can elongate MOS. Indeed, all SS isoforms can elongate MOS in vitro (Brust et al., 2013; Cuesta-Seijo et al., 2016), and may to some degree compensate for the loss of PHS1 function. Indeed, previous work in barley noted that mutants of SS1 have altered B-type granule numbers (Sparla et al., 2014). MOS can also be shortened/elongated by the disproportionating enzyme DPE1 (Critchley et al., 2001; Lütken et al., 2010; Hwang et al., 2016a), and shortened by amylolytic activities. On one hand, such redundancy in MOS processing activity suggests that multiple enzymes can likely participate in B-type granule initiation in the absence of PHS1. On the other hand, it is remarkable that loss of PHS1 alone results in such drastic reductions in B-type granule number, despite the redundancy. Clearly, PHS1 is a critical player in B-type granule initiation.

Since the initiation of A- and B-type granules is specific to the Triticeae, the role of PHS1 in B-type initiation must be limited to species within the tribe. Interestingly, RNAi suppression of PHS1 in barley led to no noticeable effects on starch synthesis (Higgins et al., 2013). It is unclear whether these findings reflect differences in the requirement of PHS1 between barley and wheat, or the incomplete silencing of PHS1 in barley. However, our results are consistent with evidence in other species that PHS1 can affect starch synthesis, although in wheat, the *phs1* phenotype appears to be restricted to granule number and less conditional on environment or other mutations. For example, rice mutants deficient in PHS1 produced grains ranging from normal to shrunken with dramatic decreases in starch content, and the shrunken grains were more prevalent when plants were grown at low temperatures (Satoh et al., 2008). In Arabidopsis, an effect on granule number is conditional upon other mutations. Complete loss of PHS1 does not affect growth, starch turnover (Zeeman et al., 2004), or granule number per chloroplast (Malinova et al., 2014) – much like in leaves of our wheat mutants (Figure 8) However, double mutants defective in PHS1 and other components of maltose metabolism have severe defects in growth and strong reductions in granule number per chloroplast (Malinova et al., 2014; Malinova and Fettke, 2017).

### Loss of PHS1 does not affect MOS levels, but affects products of MOS extension

There is substantial biochemical evidence that PHS1 can act in both biosynthetic and degradative directions *in vitro* (Kruger and ap Rees, 1983; Hwang et al., 2010; Hwang et al., 2016b) and has high affinity for soluble MOS in comparison to branched or crystalline substrates (Preiss et al., 1980; Steup and Schachtele, 1981; Steup et al., 1983). Several roles of PHS1 have been proposed, including 1) the phosphorolytic degradation of soluble glucans released during starch degradation (Steup et al., 1983), 2) the phosphorolytic degradation of linear MOS generated through amylopectin trimming during starch synthesis (Tickle et al., 2009), 3) the synthesis of MOS used for starch granule initiation (Malinova et al., 2014; Mérida and Fettke, 2021). There is currently no genetic evidence that PHS1 is required for starch degradation in any species; whereas there is genetic evidence for a conditional role in starch synthesis in *Chlamydomonas*, rice and Arabidopsis (Dauvillée et al., 2006; Satoh et al., 2008; Malinova et al., 2014). Our discovery that PHS1 is required for normal B-type granule initiation in wheat provides further support for a role of the enzyme in starch synthesis.

However, whether PHS1 fulfils its role in B-type granule initiation by acting in a biosynthetic or degradative direction is less clear. Given that elongation of MOS is thought to be a critical step in granule initiation (Nakamura, 2015; Seung and Smith, 2019; Mérida and Fettke, 2021), the simplest scenario is that PHS1 acts in the biosynthetic direction to elongate MOS released from A-type granules. Once the MOS reaches a certain length, it can act as a substrate for branching enzyme and develop more structural complexity, and initiate the formation of a B-type starch granule. However, we cannot completely rule out that PHS1 acts in the degradative direction to support granule initiation, such as trimming longer MOS to a size that is most suitable for other enzymes to act on and prime B-type starch granule initiation. Distinguishing between these two possibilities will require further research.

Our results in wheat are in contrast to recent reports that lines deficient in PHS1 have substantial MOS accumulation in rice endosperm and potato tuber (Dong et al., 2023; Flores-Castellanos and Fettke, 2023). In wheat endosperm, loss of PHS1 did not alter MOS levels (Supplemental Figures 4 and 5). Although the total MOS pool was not affected, it appears that two processes utilising MOS as a substrate were affected: granule initiation (as discussed above), and amylose synthesis. Amylose is synthesised by the Granule Bound Starch Synthase (GBSS) and is thought to be primed by MOS such as maltotriose (Denyer et al., 1996; Denyer et al., 2001; Zeeman et al., 2002; Seung, 2020). The *phs1* double mutants consistently had lower amylose content (Table 2). Interestingly, reduced amylose content was also observed in *bgc1-1*, and also in rice *flo6 (bgc1)* mutants (Peng et al., 2014; Zhang et al., 2021). It is possible that PHS1 and BGC1 influences MOS metabolism in a way that affects the priming of both B-type granules and amylose, or that the lower amylose content is a direct consequence of having low B-type granules.

### BGC1 can interact with multiple proteins

Our discovery that BGC1 interacts with PHS1 in wheat highlights yet another interaction partner for this protein, where the genetic evidence supports both partners acting in the same process (Figure 1, Table 1). In Arabidopsis, PTST2 interacts with SS4, but in the rice endosperm, it is reported to interact with ISA1, SS4 and GBSS (Peng et al., 2014; Seung et al., 2017; Zhang et al., 2021). None of these proteins were pulled down from wheat extracts, which suggests these interactions can be diverse among different species and organs. Diverse interactions could also explain how BGC1 plays multiple roles during grain development in wheat. For example, BGC1 also plays an important role in A-type granule formation during early grain development. Partial reduction in *bgc1* function reduces the number of B-type granules but permits A-type granule formation (as in the *bgc1-1* mutant), but complete elimination of BGC1 function results in the formation of compound or semi-compound granules in place of normal A-type granules (Suh et al., 2004; Chia et al., 2020). These supernumerary initiations suggest that BGC1 is involved in establishing a single initiation in amyloplasts during granule initiation. It is likely that it fulfils this role with SS4, since the *ss4* mutant also produces compound granules (Hawkins et al., 2021).

PHS1 itself has several interaction partners reported, and participates in protein complexes. In wheat and maize endosperm, PHS1 has been observed in complex with SBEI and SBEII (Tetlow et al., 2004; Subasinghe et al., 2014). Further, the rice PHS1 interacts with DPE1 (Hwang et al., 2016a). The interactions with BE and DPE1 greatly enhance the ability of PHS1 to make insoluble substrates (Nakamura et al., 2012; Hwang et al., 2016a; Nakamura et al., 2017). It is therefore of particular interest that both SBEI and DPE1 were pulled down with BGC1 (Table 1). We are currently dissecting the dynamics with which PHS1 interacts with BGC1 and these other interaction partners.

### PHS1 as a target for starch modification in the Triticeae

Since PHS1 mutations did not affect plant growth or grain yield (Figure 3), PHS1 represents another gene target to reduce the content of B-type granule content in wheat. Varieties with low B-type granules are desirable for bread making (Park et al., 2009). Recent work with BGC1 mutants showed that starch without B-type granules had higher water absorption, reduced grain hardness and higher protein content (Saccomanno et al., 2022). B-type granules also cause processing problems due to their small size (Stoddard and Sarker, 2000; Park et al., 2009). Further, reduced B-type granules are highly desired in barley for brewing, since B-type granules cause filtration problems in brewing and cause a starch haze (Stark and Lynn, 1992). Since reduced B-type granules can only be achieved through BGC1 by reducing gene dosage rather than through knockouts, the phenotype is easier to achieve in a polyploid species like wheat than in a diploid species like barley. Thus, PHS1 could be a more suitable gene target for reducing B-type granules in diploid Triticeae as the phenotype can be achieved through a homozygous knockout mutation.

## Materials and Methods

### Plant materials and growth

Wheat plants were grown in climate-controlled glasshouses for all grain analyses, and in controlled environment rooms (CERs) for the analysis of starch in leaves. The glasshouses were set to provide a minimum 16 h of light at 20°C and 16°C during the dark. The CERs were set to provide 16 h light at 20°C (light intensity was 300 μmol photons m^−2^ s^−1^) and 8 h dark at 16°C. *N. benthamiana* were grown in climate-controlled glasshouses set to provide a minimum of 16 h light (200 μmol photons m^−2^ s^−1^) and a constant temperature (22°C). All glasshouses and CERs were set to provide 60% relative humidity.

Mutant lines of *PHS1* in durum wheat (*Triticum turgidum* cv. Kronos) were obtained from the wheat *in silico* TILLING resource (http://www.wheat-tilling.com) (Krasileva et al., 2017). Lines Kronos4533(K4533) and Kronos4367(K4367) were obtained for *PHS1-5A*, and lines Kronos2864(K2864) and Kronos0238(K0238) were obtained for *PHS1-5B* (Figure 2). Plants were crossed to combine mutations in the A- and B-homoeologs. The wild-type segregant (AA BB), single homeolog mutants (*aa* BB, AA *bb*) and the double homeolog mutant (*aa bb*) were selected in the F2 generation using KASP V4.0 genotyping (LGC, Teddington) with the primers in Supplemental Table 1. The *bgc1-1* mutant in Kronos is previously described in Chia et al. (2020).

### Isolation, cloning, plasmid construction for PHS1 and BGC1

Total RNA was extracted from leaves of two-week-old wheat seedlings using RNease kit (Qiagen) with on-column DNase I digestion (Qiagen). cDNA was synthesized (using 2 µg RNA) using the Go Script™ Reverse Transcriptase Kit (Promega) following the manufacturer’s instruction. The full-length wheat *PHS1* cDNA sequence was amplified using gene-specific primers listed in Supplemental Table 2 and inserted into Gateway entry vector, pENTR, using the pENTR/D-TOPO kit (Invitrogen). To produce constructs with a C-terminal YFP or RFP tag, we recombined the coding sequences from PHS1:pENTR and BGC1:pDONR221 (Hawkins et al., 2020) into the plant expression vectors pB7YWG2 (35S promoter; C-terminal YFP) or pB7RWG2 (35S promoter; C-terminal RFP) using Gateway LR clonase (Invitrogen). To generate the chloroplast-targeted YFP and RFP proteins, the rubisco small subunit (RbcS) transit peptide (Kim et al., 2010) was cloned into pDONR221, and recombined into the pB7YWG2 and pB7RWG2.

### Transient expression of PHS1 and BGC1 in N. benthamiana and protein localisation

Constructs for YFP-tagged PHS1 and BGC1 expression were transformed into *Agrobacterium tumefaciens* (strain GV3101). *Agrobacterium* cells were grown at 28°C in Luria-Bertani medium with the appropriate antibiotics. Cells were pelleted, washed, and resuspended in the infiltration medium [10 mM 2-(N-morpholino) ethanesulfonic acid, 10 mM MgCl2, and 0.15 mM acetosyringone, pH 5.6]. The cell suspension was then infiltrated into the intercellular spaces between abaxial epidermal cells of intact *Nicotiana benthamiana* leaves using a 1 mL plastic syringe as previously described (Kamble and Majee, 2022). Infiltrated plants were incubated overnight in dark followed by 2 days in a 16 h/8 h light–dark cycle. YFP fluorescence in leaf samples was observed using a laser scanning confocal microscope (SP8; Leica) with a 63× water-immersion lens. Images were processed using LAS X software.

### Pulldown assay, immunoprecipitations and mass spectrometry

For the pulldown assay to identify proteins associating with BGC1, the bait protein (recombinant His-tagged BGC1) was expressed in *E. coli* as described in Hawkins et al. (2020) and purified in its native state using Ni-NTA agarose beads (Qiagen) as previously described (Seung et al., 2013). To produce endosperm extract, endosperms were dissected from developing grains (collected at 18 dpa) of wild-type Kronos plants and homogenised in ice-cold extraction medium [50 mM Tris-HCl, pH 8, 1 mM DTT, 1% (v/v) Triton X-100, 150 mM NaCl, and Roche Complete Protease Inhibitor cocktail] at a rate of 1 mL buffer per 100 mg tissue. Insoluble material was removed by centrifugation at full speed for 5 min at 4°C, and proteins were collected in the supernatant. Recombinant BGC1-His protein (2.5 µg) was added to the supernatant (1 mL) and was incubated for 1 h at 4°C. µMACS magnetic beads conjugated to anti-His (Miltenyi Biotec) were added and incubated for 1 h at 4°C to retrieve the bait protein together with interacting proteins. The beads were captured with a µColumn on a magnetic stand (Miltenyi Biotec), washed three times with wash medium [50 mM Tris-HCl, pH 8, 1 mM DTT, 1% (v/v) Triton X-100, 300 mM NaCl, and Roche Complete Protease Inhibitor cocktail], then three times with wash medium without Triton X-100, before eluting the bound proteins with elution medium [50 mM Tris-HCl, pH 6.8, and 2% (w/v) SDS].

The eluted proteins were precipitated with chloroform/methanol (Pankow et al., 2016). Protein pellets were resuspended in 50 µl of 2.5% sodium deoxycholate (SDC; Merck) in 0.2 M EPPS-buffer (Merck), pH 8.5 and reduced, alkylated, and digested with trypsin in the SDC buffer according to standard procedures. After the digest, the SDC was precipitated by adjusting to 0.2% trifluoroacetic acid (TFA), and the clear supernatant subjected to C18 SPE. Samples were dried in a SpeedVac concentrator (Thermo Fisher Scientific, #SPD120) and the peptides dissolved in 0.1%TFA/3% acetonitrile.

Peptides were analysed by nanoLC-MS/MS on an Orbitrap Eclipse™ Tribrid™ mass spectrometer with a FAIMS Pro Duo source, coupled to an UltiMate® 3000 RSLCnano LC system (Thermo Fisher Scientific, Hemel Hempstead, UK). The samples were loaded and trapped using a trap cartridge (Pepmap Neo C18, 5 µm, 300 µm x 5 mm, Thermo) with 0.1% TFA at 15 µl min^-1^ for 3 min. The trap was then switched in-line with the analytical column (nanoEase M/Z column, HSS C18 T3, 100 Å, 1.8 µm; Waters, Wilmslow, UK) for separation using the following gradient of solvents A (water, 0.1% formic acid) and B (80% acetonitrile, 0.1% formic acid) at a flow rate of 0.2 µl min^-1^ : 0-3 min 3% B (during trapping); 3-10 min linear increase B to 7%; 10-100 min increase B to 32%; 100-148 min increase B to 50%; followed by a ramp to 99% B and re-equilibration to 3% B, for a total running time of 180 minutes. Mass spectrometry data were acquired with the FAIMS device set to three compensation voltages (−35V, −50V, −65V) at standard resolution for 1 s each with the following MS settings in positive ion mode: MS1/OT: resolution 120K, profile mode, mass range m/z 300-1800, spray voltage 2800 V, AGC 4e^5^, maximum injection time 50 ms; MS2/IT: data dependent analysis was performed using HCD fragmentation with the following parameters: cycle time of 1 s in IT turbo for each FAIMS CV, centroid mode, isolation window 1.0 Da, charge states 2-5, threshold 1.0e^4^, CE = 30, normalised AGC target 100%, max. inject time set to Auto, dynamic exclusion 1 count, 15 s exclusion, exclusion mass window ±10 ppm. The acquired raw data were processed and quantified in Proteome Discoverer 3.0 (Thermo) using the incorporated search engine CHIMERYS® (MSAID® Munich, Germany). The processing workflow included recalibration of MS1 spectra (RC) and the Minora Feature Detector for quantification with min. trace length=7 and S/N threshold=3. The Top N Peak Filter (10 per 100 Da) was applied and the CHIMERYS® search was performed with the prediction model inferys_2.1_fragmentation, enzyme trypsin with 2 missed cleavages, peptide length 7-25, fragment tolerance 0.5 Da, variable modification oxidation (M), fixed modifications carbamidomethyl (C). Percolator was used for validation using q-value and FDR 0.01 (strict) and 0.05 (relaxed).

In the consensus workflow quantification was performed with a maximum RT Shift of 3 min and a mass tolerance of 4 ppm between runs. Protein quantification was based on the top 3 most abundant unique peptides per protein group. Missing values were replaced by low abundance resampling. Protein abundance ratios were calculated from the 3 replicates per sample. The hypothesis test was performed by a background-based t-test and the p-values adjusted according to BH.

For pairwise co-immunoprecipitations, proteins were transiently expressed in *N. benthamiana* leaves as described above, and proteins were extracted as described for the pulldown assay. The supernatant was incubated for 1 h at 4°C with µMACS magnetic beads conjugated to anti-YFP. After incubation, the beads were recovered using a µColumn (Miltenyi Biotec) on a magnetic stand. The beads were washed five times with wash medium [50 mM Tris-HCl, pH 8, 1 mM DTT, 1% (v/v) Triton X-100, 300 mM NaCl, and Roche Complete Protease Inhibitor cocktail] before eluting the bound proteins with SDS-PAGE loading buffer [50 mM Tris-HCl, pH 6.8, 2% (w/v) SDS, 100 mM DTT, 3% (v/v) glycerol, 0.005% (w/v) bromophenol blue]. The eluates were analysed using SDS-PAGE and immunoblotting, using anti-YFP (Torrey Pines; TP401 - 1:5000) and anti-RFP (Abcam plc; ab34771 - 1:2000) primary antibodies. Proteins were detected using chemiluminescence from horseradish peroxidase-coupled secondary antibodies Anti-rabbit HRP (Sigma; A0545), 1:15,000.

### Native gel analysis of PHS activity

The visualisation of ⍺-glucan phosphorylase activity by native PAGE was carried out using the method of (Zeeman et al., 2004) with minor modifications. Leaf samples were homogenised in extraction medium [100 mM MOPS (3-(N-morpholino) propane sulfonic acid), pH 7.2, 1 mM DTT, 1 mM EDTA, 10% (v/v) ethanediol, and Complete protease inhibitor cocktail (Roche)], then spun at 20,000*g* for 10 min. Soluble proteins were collected in the supernatant, and the protein concentration of the extracts was determined using the Bradford protein assay. Equal amounts of protein (∼100-180 µg) were loaded per lane on native PAGE gels (7.5% acrylamide and 0.3% oyster glycogen in resolving gel; 3.75% acrylamide in stacking gel) and were electrophoresed at 100V for 4 h at 4°C. Gels were washed twice in 100 mM Tris-HCl, pH 7.0, 1 mM DTT, then incubated in 100 mM Tris-HCl, pH 7.0, 1 mM DTT, 50 mM Glc-1-P overnight at 20°C. Activity bands were stained with Lugol’s iodine solution (L6146, Sigma, St. Louis).

### Grain morphometrics

Grain yield per plant was quantified as the total weight of grains harvested from each plant. The thousand grain weight and grain size were quantified using the MARViN seed analyser (Marvitech GmbH, Wittenburg).

### Starch content of leaves and mature grains

Starch content of leaves and mature grains (in glucose equivalents) were quantified as described in Hawkins et al. (2020). Briefly, for leaves, two-week-old seedlings were harvested at the base of the lowest leaf and flash frozen in liquid N_2_, then homogenised in 0.7 M perchloric acid. Insoluble material was collected by centrifugation, washed three times in 80% ethanol, then resuspended in water. Starch was digested using α-amylase/amyloglucosidase (Roche, Basel), and the released glucose was assayed using the hexokinase/glucose-6-phosphate dehydrogenase assay (Roche). For mature grains, flour (5-10 mg) was dispersed in 100 mM sodium acetate buffer, pH 5, and the starch was digested with thermostable α-amylase at 99°C for 7 min. Amyloglucosidase was added to the digestion, and was further incubated at 50°C for 35 min. Both the thermostable α-amylase and amyloglucosidase were from the Total Starch Assay kit (K-TSTA, Megazyme, Bray). The digested sample was centrifuged to remove insoluble material, and glucose was measured in the supernatant as for the leaf starch quantification.

### Starch purification and determination of granule morphology, size distribution and amylose content

For starch purification from mature grains, grains (3-5 grains per extraction) were soaked overnight in ddH_2_O at 4°C, then homogenised in a mortar and pestle with excess ddH_2_O. The homogenates were filtered through a 70 µm nylon mesh, then centrifuged at 3000*g*, 5 min, before resuspending the starch pellet in water. The starch suspension was centrifuged at 2500*g*, 5 min on a cushion of 90% (v/v) Percoll, 50 mM Tris-HCl, pH 8. The pellet was washed twice in 50 mM Tris-HCl, pH 6.8, 10 mM EDTA, 4% SDS (v/v), 10 mM DTT, then twice in ddH_2_O, before finally resuspending in ddH_2_O.

The morphology of purified granules was examined using a Nova NanoSEM 450 (FEI, Hillsboro) scanning electron microscope (SEM). To quantify starch granule size distributions, the purified starch was suspended in Isoton II electrolyte solution (Beckman Coulter, Indianapolis), and particle sizes were measured using a Multisizer 4e Coulter counter (Beckman Coulter) fitted with a 70 µm aperture tube. A minimum of 100,000 particles was measured per sample. These data were used to produce relative volume vs. diameter and relative number vs. diameter plots. A mixed bimodal (two log-normal distributions) were fitted to the relative volume vs. diameter plots to calculate mean diameters of A- and B-type granules, and the B-type granule volume percentage (defined as volume occupied by B-type granules as a percentage of the total volume of starch).

Amylose content of the purified starch granules was estimated using an iodine colourimetry method (Washington et al., 2000). Briefly, starch (1 mg) was dispersed overnight at room temperature in 1 M NaOH. The solution was neutralised to pH 7 using 1 M HCl, and 5 µL of this solution was diluted in 220 µL of water and 25 µL Lugol’s iodine solution (L6146, Sigma, St. Louis). The reaction was incubated at room temperature for 10 minutes prior to absorbance measurements at 535 nm and 620 nm. Apparent amylose content was estimated using the formula:

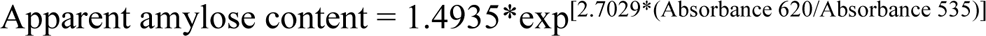

### Quantification of starch, sugars, MOS, granule number and size in developing grains

Endosperms were dissected from developing grains collected at 8, 14, 18 and 22 dpa. In each extraction, three to five individual endosperms of known fresh weight were used. After homogenisation in 0.7 M perchloric acid, homogenates were spun at 10,000g for 5 mins, and the supernatant was immediately neutralized using neutralisation buffer (2 M KOH, 400 mM MES). This neutralised soluble fraction was used for sugar and MOS quantification (see below).

The pellet was resuspended in ddH_2_O and equally divided into two fractions. One fraction was used to quantify starch content following the method described above for leaves. The other was used for starch purifications (as described above for mature grains), and all granules purified within this fraction was resuspended in known volumes of Isoton II electrolyte solution (Beckman Coulter, Indianapolis). The suspension was analysed in a Multisizer 4e Coulter counter (Beckman Coulter) fitted with a 70 µm aperture tube and running in volumetric mode (analysing 2 mL of the suspension). This gave the number of granules in the suspension, which could be used to calculate the number of granules per starting fresh weight of endosperm; as well as granule size distribution plots.

For the quantification of soluble glucans: Total soluble glucans (neutralised soluble fraction without precipitation) and methanol-precipitable soluble glucans (after precipitation) were quantified. For methanol precipitation, one volume of neutralised soluble fraction was mixed with 4 volumes of pure methanol, mixed, and incubated overnight at −20°C. Precipitated glucans were collected by centrifugation at 10,000*g*, 5 min, then washed with 75% methanol, and dried. Dry pellets were suspended in one volume of ddH_2_O. To quantify glucans, the neutralised soluble fraction and resuspended precipitated glucans were digested using α-amylase/amyloglucosidase (Roche, Basel), and released glucose was assayed using the hexokinase/glucose-6-phosphate dehydrogenase assay (Roche).

HPAEC-PAD was used to quantify glucose, fructose, sucrose, and maltose, and to visualise MOS accumulation patterns. The neutralised soluble fraction was purified on sequential columns of Dowex 50W and Dowex 1 (Sigma) as described previously Seung et al. (2013). Purified samples were separated on an ICS-5000 HPLC fitted with a CarboPac^TM^ PA20 column (3 × 250 mm, CV=1.06 ml; Dionex). The mobile phase consisted of eluate A (100 mM NaOH) and eluate B (150 mM NaOH, 500 mM sodium acetate), following a gradient program of: 0–7 min, 0% B; 7.0–26.5 min, a concave gradient to 80% B; 26.5–32.0 min, 80% B; 32.0-32.1 min, linear gradient to 0% B, 32.1–40.0 min, 0% B with flow rate of 0.25 ml/min.

### Light microscopy of leaf sections

Starch granules were visualised in leaf chloroplasts as described in Chen et al. (2023). Briefly, 1.5 mm x 1.5 mm squares leaf samples were harvested at the base of the lowest leaf from two-week-old seedlings in fixative [glutaraldehyde (2.5%), sodium cacodylate, pH 7.4 (0.05 M)]. Samples were then post-fixed with osmium tetroxide (1% w/v) in sodium cacodylate, pH 7.4 (0.05 M), After dehydration in an ascending series of ethanol, samples were embedded in LR white resin using EM TP embedding machine (Leica). Semi-thin sections (0.5 µm thick) were produced from the embedded leaves using a glass knife and were dried onto PTFE-coated slides. Starch was stained using reagents from the Periodic Acid-Schiff staining kit (Abcam): using a 30 min incubation with periodic acid solution, followed by 5 min with Schiff’s solution. Chloroplasts and cell walls were stained using toluidine blue stain (0.5% toluidine blue ‘O’, 0.5% sodium borate) for 1 min. The sections were mounted with Histomount (National Diagnostics) and imaged on a DM6000 microscope with 63X oil immersion lens (Leica).

### Statistical analysis

Statistical analyses were carried out using the SPSS program (SPSS Statistics, IBM).

### Accession numbers

The accession numbers corresponding to the genes investigated in this study are: *TraesCS5A02G395200 (PHS1-A1), TraesCS5B02G400000 (PHS1-B1), TRITD5Av1G205670 (PHS1-A1), TRITD5Bv1G201740 (PHS1-B1), TraesCS4A02G284000 (BGC1-A1), TraesCS4B02G029700 (BGC1-B1), TRITD4Av1G198830 (BGC1-A1), TRITD0Uv1G034540 (BGC1-B1)*.

### Data Availability

The mass spectrometry proteomics data have been deposited to the ProteomeXchange Consortium via the PRIDE partner repository with the dataset identifier PXD042396 and Project DOI:10.6019/PXD042396.

## Acknowledgements

We thank Kay Trafford and Tansy Chia (NIAB) for providing *bgc1-1* seeds, Elaine Barclay (John Innes Centre - JIC) for assistance with leaf sectioning, and Martin Rejzek (JIC) for advice with HPAEC-PAD analysis. We thank JIC Horticultural Services for providing growth facilities and maintenance of plant material, JIC Proteomics for providing access to mass spectrometry, JIC Bioimaging for providing access to microscopes, JIC Genotyping for providing support for KASP genotyping and JIC Chemistry for providing access to HPAEC-PAD instruments. This work was funded through a John Innes Foundation (JIF) Chris J. Leaver Fellowship (to D.S), a Biotechnology and Biological Sciences Research Council (BBSRC, UK) research grant BB/W015935/1 (to D.S.), the BBSRC-funded John Innes Centre Flexible Talent Mobility Account (BB/S507921/1), and the BBSRC-funded Institute Strategic Programme Harnessing Biosynthesis for Sustainable Food and Health (HBio) (grant number BB/X01097X/1).

## Author contributions

N.U.K. and D.S. conceived and designed the study. All authors performed the research and analysed data. N.U.K and D.S. wrote the article with input from all authors.

## Supplemental Figures and Tables

**Supplemental Figure 1**: Expression levels of PHS1 homoeologs in different tissues of bread wheat.

**Supplemental Figure 2:** Size distribution of grains from *phs1-1* mutants.

**Supplemental Figure 3**: Endosperm starch from the *phs1-6* mutant has fewer and larger B-type granules.

**Supplemental Figure 4:** Soluble glucan quantification in the developing endosperm tissue.

**Supplemental Figure 5:** Soluble sugar quantification in the developing endosperm tissue.

**Supplemental Figure 6:** Starch granules in leaf chloroplasts of the *phs1-6* mutant.

**Supplemental Data 1:** List of BGC1 interactors identified using mass spectrometry.

**Supplemental Table 1:** Primers used for KASP genotyping.

**Supplemental Table 2:** Oligonucleotides used in this study.

