## Supplemental Figures and Tables for "Initiation of B-type starch granules in wheat endosperm requires the plastidial α-glucan phosphorylase PHS1"

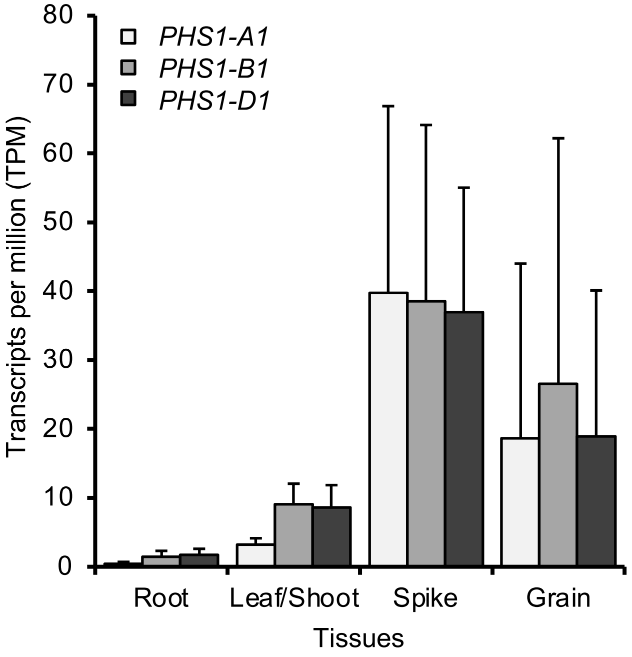


**Supplemental Figure 1**: Expression levels of PHS1 homoeologs in different tissues of bread wheat. Data are from cultivar Chinese Spring, and obtained using the wheat expression browser (http://www.wheat-expression.com/; Borrill et al. 2016).

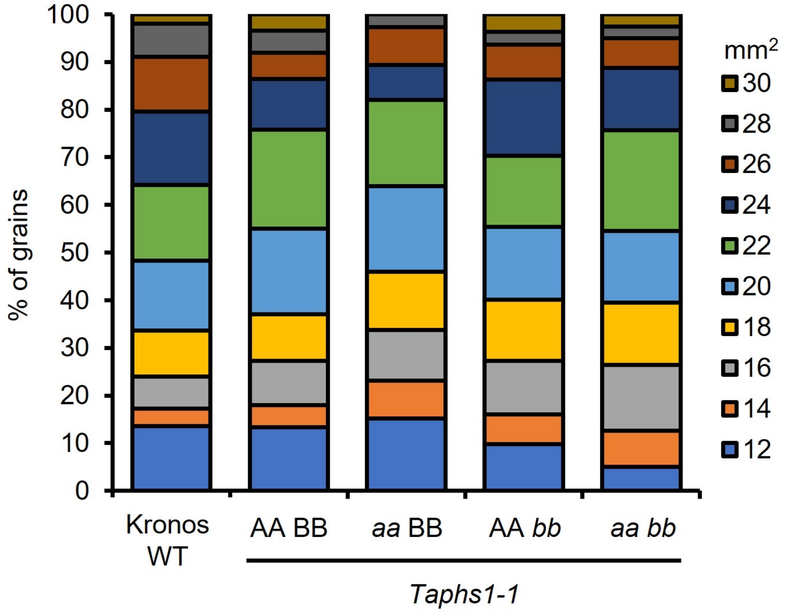


**Supplemental Figure 2**: Size distribution of grains from *phs1-1* mutants. Grain size data (calculated based on 2-D area) are from the mature grains. Grains from $n = 5$ plants per genotype were pooled for analysing the size distribution (representing a total of 167-368 individual grains per genotype).

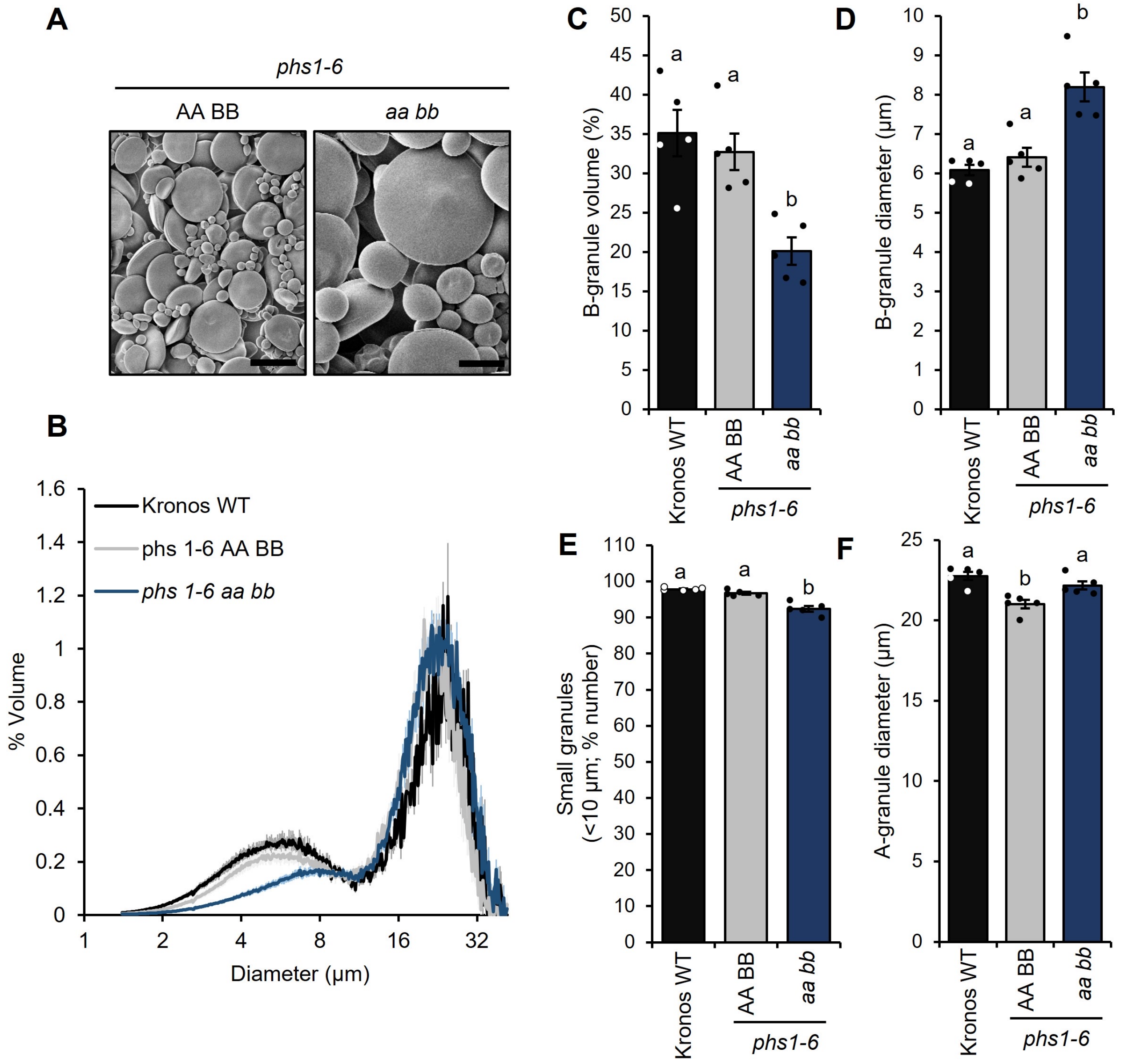


**Supplemental Figure 3**: **Endosperm starch from the *phs1-6* mutant has fewer and larger B-type granules. A**) Scanning electron micrographs of purified endosperm starch. Bar = 10 µm. **B**) Granule size distributions were determined using a Coulter counter, and the data were expressed as relative % volume (of total starch) vs. granule diameter plots. **C** and **D**) B-type granule volume (% of total starch) and the average diameter of B-type granules were extracted from the relative volume vs. diameter plots by fitting a bimodal mixed normal distribution. **E**) The percentage of small granules by number (smaller than 10 µm) were calculated from the coulter counter data. **F**) The average diameter of A-type granules was calculated from the relative volume vs. diameter plots, as for panels C and D. For panels B-F, plots show the mean from the analysis of n=5 replicate starch extractions, each from grains from a separate plant. The shading (on panels B) and error bars (on panels C, D, E, F) represent the SEM. Values with different letters are significantly different under a one-way ANOVA with Tukey's post-hoc test ($p < 0.05$).

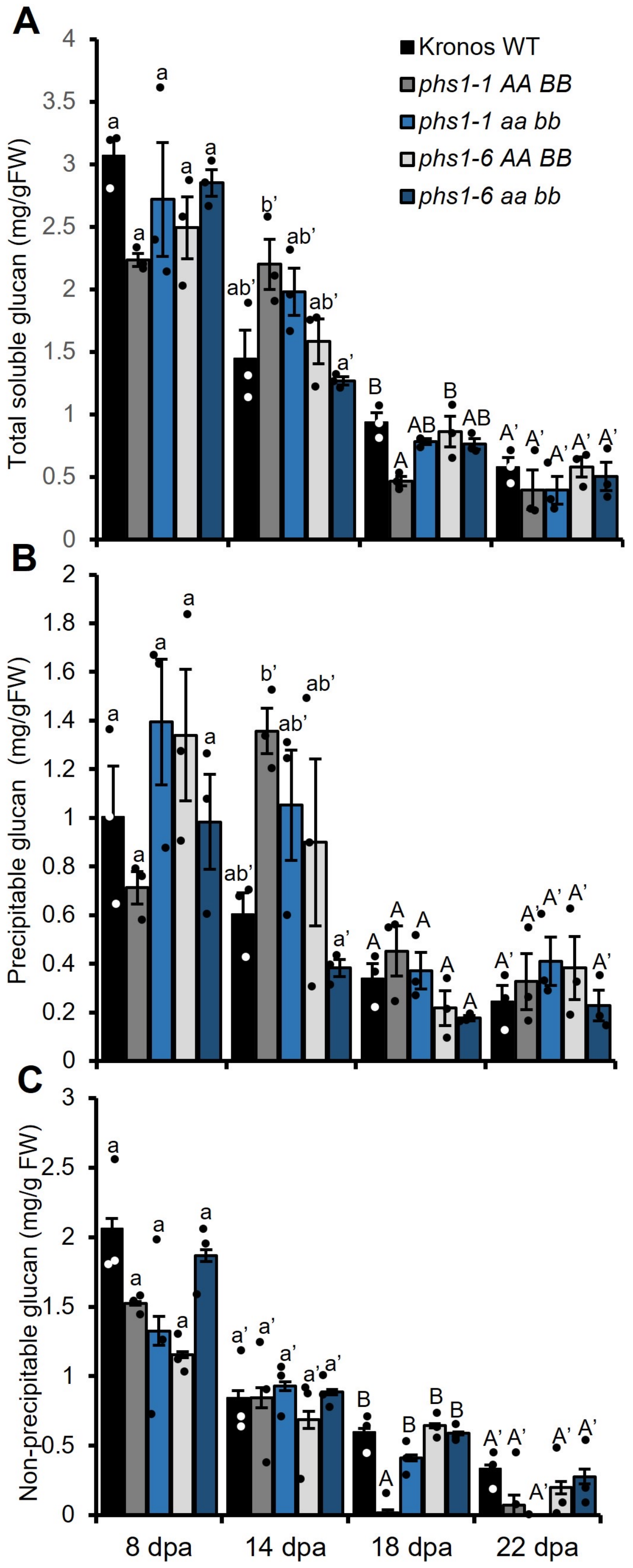


**Supplemental Figure 4: Soluble glucan quantification in the developing endosperm tissue. A)** Total soluble glucans, **B**) methanol precipitable fraction of soluble glucans (phytoglycogen and long MOS), and **C**) the non-precipitable fraction (short MOS) in perchloric acid extracts of grain of WT, *phs1-1* and *phs1-6* double mutants and corresponding wild-type controls, harvested at 8, 14, 18 and 22 days post anthesis (dpa), with n = 3 individual plants for each genotype per time point. Values are expressed relative to the fresh weight of the dissected endosperm. Values with different letters are significantly different under a one-way ANOVA with Tukey's post-hoc test ($p < 0.05$).

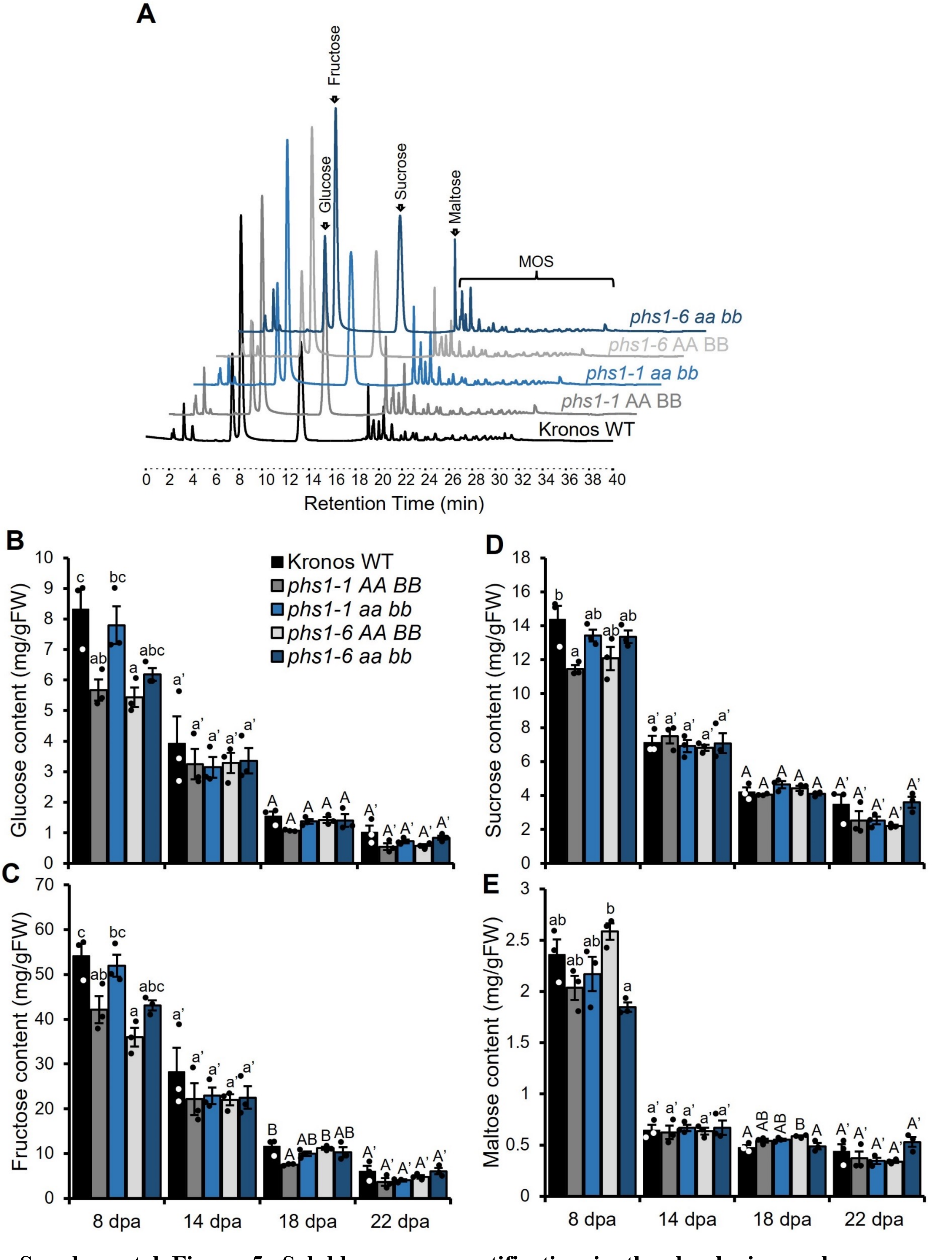


**Supplemental Figure 5**: **Soluble sugar quantification in the developing endosperm tissue. A**) Representative HPAEC-PAD chromatograms, used for the quantification of **B**) glucose, **C**) fructose, **D**) sucrose and **E**) maltose. Perchloric acid extracts of dissected endosperm harvested at 8, 14, 18 and 22 days post anthesis (dpa) were analysed, with $n = 3$ individual plants for each genotype per time point. Values are expressed relative to the fresh weight of the dissected endosperm. Values with different letters are significantly different under a one-way ANOVA with Tukey's post-hoc test ($p < 0.05$).

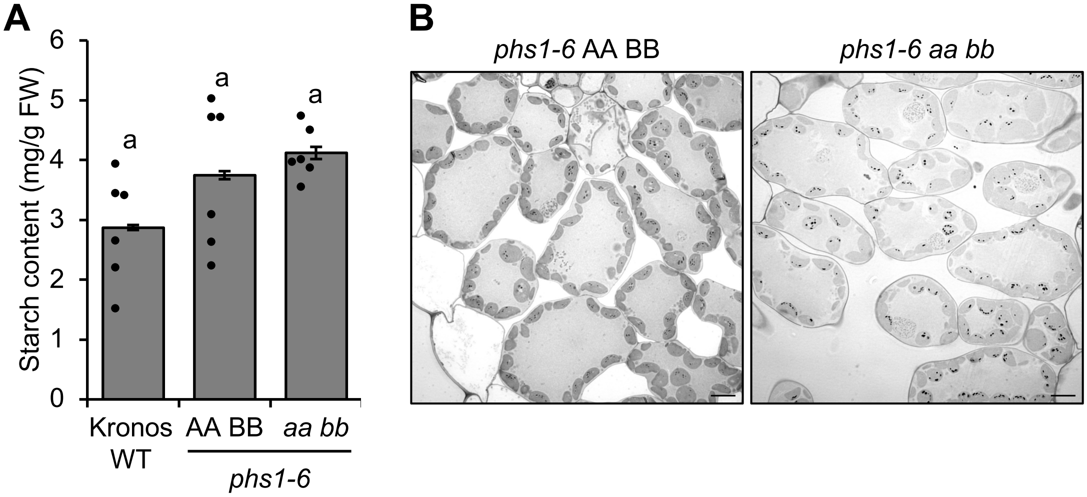


**Supplemental Figure 6: Starch granules in leaf chloroplasts of the *phs1-6* mutant**. **A)** Total starch content of leaves of the Kronos WT, wild-type segregant control (AA BB) and double mutant (*aa bb*). Seedlings were grown for two weeks under 16 h day/8 h night and harvested at the end of day. Values are the mean±SEM from n = 6 plants (each represented by a data point). There were no significant differences under a one-way ANOVA and Tukey's posthoc test at $p < 0.05$. **B)** Light micrographs of leaf sections stained with Periodic Acid-Schiff (PAS) staining to visualise starch granules. Leaf segments were harvested halfway along the length of the older blade in two-week-old seedlings were harvested at the ED, prior to fixation and embedding.

| **Supplemental Table 1:**<br>Primers used for KASP genotyping | | | |
|---|---|---|---|
| Allele | WT | mutant | common |
| K4533 | tttttgattctctgatattgaattg | tttttgattctctgatattgaatta | aaggtatgagataaggctgaaga |
| K4367 | atatccgtttcgacctgcag | atatccgtttcgacctgcaa | ccatatgtccttggcatactcg |
| K2864 | agagacatcatttcttacgatctcc | agagacatcatttcttacgatctct | acctgcatgttagcttcttttct |
| K0238 | aaaataaagtagacgaaagaatac | aaaataaagtagacgaaagaatat | acaaaattttgatttgggtgcc |
| K2244 | ttgtcaagagaccatgttcgc | ttgtcaagagaccatgttcgt | ctcctagtcgaagtccaacta |
| K3239 | gcccaatcctgcttcagaag | gcccaatcctgcttcagaaa | gccacccattttgtagttagttag |
| All primer sequences are provided in 5' to. 3' orientation. For KASP genotyping, the VIC/HEX tail (gaaggtcggagtcaacggatt) was added to the 5' end of the WT allele primers, and the FAM tail (gaaggtgaccaagttcatgct) was added to the 5' end of the mutant allele primers. | | | |

| **Supplemental Table 2:**<br>Oligonucleotides used in this study.<br><br>Cloning of PHS1:pENTR | |
|---|---|
| Forward primer | Reverse primer |
| CACCATGGCGACCGCCTCGCCGCCGCTCG | GGGCATGATGACGGGGCTGATACC |
